# Direct current stimulation boosts Hebbian plasticity in vitro

**DOI:** 10.1101/562322

**Authors:** Greg Kronberg, Asif Rahman, Mahima Sharma, Marom Bikson, Lucas C. Parra

## Abstract

**Background:** There is evidence that transcranial direct current stimulation (tDCS) can improve learning performance. Arguably, this effect is related to long term potentiation (LTP), but the precise biophysical mechanisms remain unknown.

**Hypothesis:** We propose that direct current stimulation (DCS) causes small changes in postsynaptic membrane potential during ongoing endogenous synaptic activity. The altered voltage dynamics in the postsynaptic neuron then modify synaptic strength via the machinery of endogenous voltage-dependent Hebbian plasticity. This hypothesis predicts that DCS should exhibit Hebbian properties, namely pathway specificity and associativity.

**Methods:** We studied the effects of DCS applied during the induction of LTP in the CA1 region of rat hippocampal slices and using a biophysical computational model.

**Results:** DCS enhanced LTP, but only at synapses that were undergoing plasticity, confirming that DCS respects Hebbian pathway specificity. When different synaptic pathways cooperated to produce LTP, DCS enhanced this cooperation, boosting Hebbian associativity. Further slice experiments and computer simulations support a model where polarization of postsynaptic pyramidal neurons drives these plasticity effects through endogenous Hebbian mechanisms. The model is able to reconcile several experimental results by capturing the complex interaction between the induced electric field, neuron morphology, and endogenous neural activity.

**Conclusions:** These results suggest that tDCS can enhance associative learning. We propose that clinical tDCS should be applied during tasks that induce Hebbian plasticity to harness this phenomenon, and that the effects should be task specific through their interaction with endogenous plasticity mechanisms. Models that incorporate brain state and plasticity mechanisms may help to improve prediction of tDCS outcomes.

## Introduction

Transcranial direct current stimulation (tDCS) studies in humans have recently exploded in number and scope (1–4). While these studies have seen varying degrees of success (1), in aggregate they suggest that stimulation with weak constant current can have long term effects on cognitive function (5). One of the predominant theories to explain these long term effects is that stimulation affects synaptic plasticity (6), although a variety of alternatives have also been proposed (7,8) and are being explored (9). The synaptic plasticity theory is consistent with an array of findings from pharmacological studies in humans (10) as well as animal electrophysiology studies conducted in-vivo (9,11,12) and in-vitro (13–16). However, the biophysical mechanism for such plasticity effects is unknown.

Polarization of neuronal membranes in response to extracellular electric fields has been well characterized (17–23), as has the membrane potential-dependence of Hebbian plasticity (24–27). While it is straightforward to draw a connection between these phenomena, their interaction can be complex. For example, we previously observed that the effects of DCS depend on both the location of active synapses and the precise temporal patterns of activity used to induce plasticity (14). These results suggest that the effects of DCS depend on the interaction between the induced electric field, neuron morphology, and the endogenous brain dynamics. Given this complexity, it is perhaps no surprise that results from human clinical trials with tDCS have remained inconclusive (28–31), or that optimization of tDCS protocols has been slow. For example, there is an ongoing debate as to whether tDCS should be applied before, during, or after a behavioral or cognitive task (32–34).

We propose that DCS causes small changes in postsynaptic membrane potential during ongoing endogenous synaptic activity. The altered voltage dynamics in the postsynaptic neuron then modify synaptic strength via the machinery of endogenous voltage-dependent Hebbian plasticity. An implication of this hypothesis is that the effects of DCS should exhibit similar properties as the endogenous Hebbian plasticity that it is paired with. Two of these properties, pathway specificity and pathway associativity (19,20), support functionally specific learning of cell assemblies in neural networks (21,22). tDCS may therefore enhance functionally specific learning by acting through this Hebbian mechanism.

We induced LTP in hippocampal brain slices using theta rhythms (theta burst stimulation, TBS), and confirm that this form of “endogenous” plasticity is pathway specific and associative. Applying DCS during plasticity induction boosted the amount of LTP, while maintaining the pathway-specific and associative properties of the underlying endogenous plasticity. Additional experiments and computer simulations support the hypothesized model in which DCS achieves these effects through altered neuronal excitability and subthreshold depolarization in dendrites during ongoing synaptic input.

We present what is, to our knowledge, the first computational model of the effects of DCS on synaptic plasticity, which reconciles several experimental results. The model makes specific and testable predictions for both how tDCS should alter plasticity when paired with various endogenous brain states, and how this can inform the design of tDCS protocols. Specifically, the most effective tDCS interventions should be those that pair stimulation concurrently with behavioral training and that performance gains should be specific to the learned task.

## Results

### Anodal DCS boosts LTP

To mimic learning during a training task we induced LTP by applying TBS in the hippocampal Schaffer collateral pathway (4 pulses at 100 Hz repeated for 15 bursts at 5 Hz, 3 seconds total). We applied acute anodal or cathodal DCS (see Methods) for the duration of the LTP-induction protocol (20 V/m; Fig. 1A). When paired with anodal DCS, the resulting LTP was increased compared to TBS alone (Figure 1B; control: 1.287+−0.025, N=52 slices; anodal: 1.397+−0.047, N=32 slices, p=0.027). However, cathodal stimulation had no significant effect (Figure 1B; cathodal: 1.243+−0.031, N=12 slices, p=0.424).

**Figure 1.**
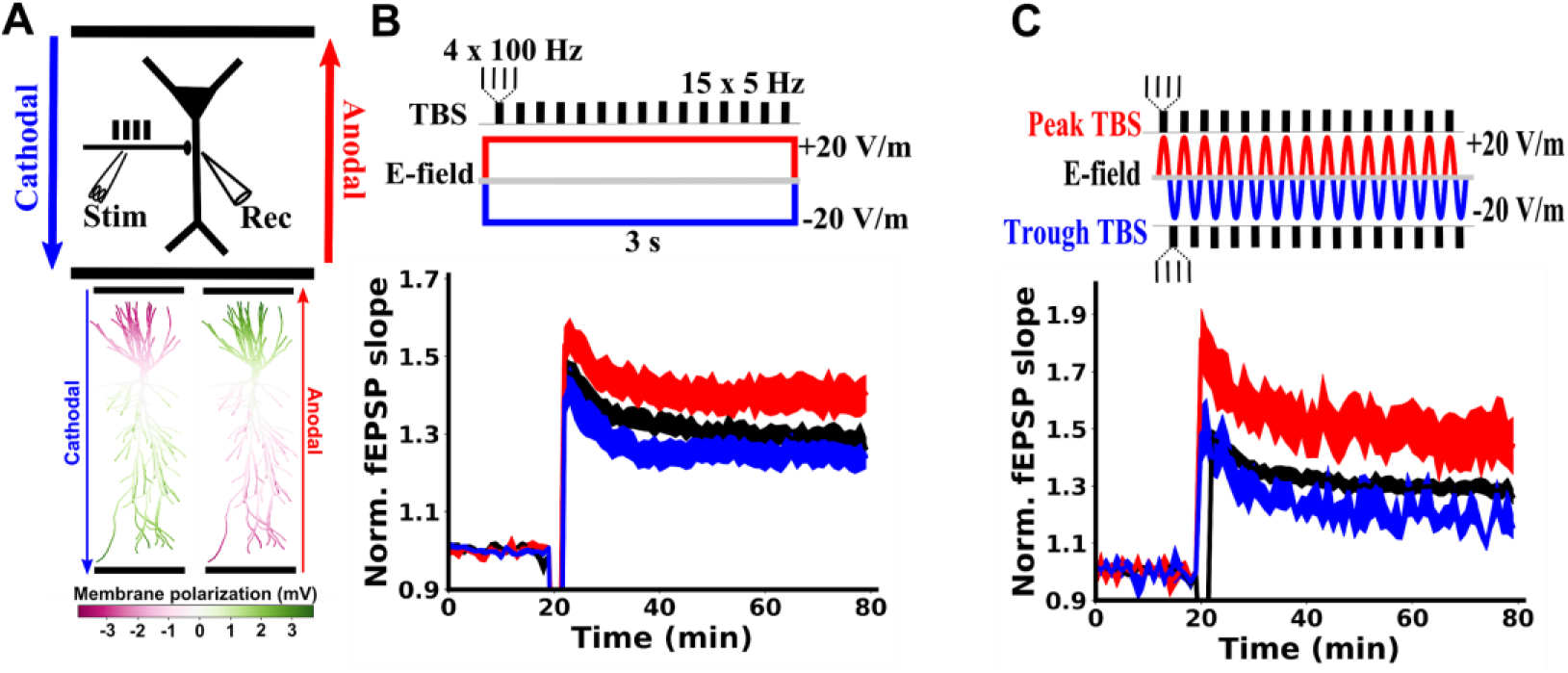
Soma-depolarizing electric fields enhance TBS-induced LTP in hippocampal Schaffer Collateral pathway. **A) Top:** Schematic of the experimental setup, showing the orientation of anodal (red) and cathodal (blue) electric fields generated by parallel wires (black horizontal lines). Location of stimulation (Stim) with TBS and recording (Rec) of field excitatory postsynaptic potentials (fEPSP) are indicated relative to a CA1 pyramidal neuron soma (black triangle). **Bottom:** Membrane polarization throughout a model pyramidal neuron in response to 20 V/m anodal (red) or cathodal (blue) DCS. Green compartments are depolarized due to DCS, while magenta compartments are hyperpolarized by DCS. **B)** Constant current stimulation applied during TBS modulates the resulting LTP measured as a change in fEPSP normalized to baseline. **C)** Alternating current stimulation (5Hz) was applied and TBS bursts were timed to either the peak (red) or the trough (blue) of the sinusoidal alternating current. Note that the applied electric field at the peak of the alternating current is identical to anodal constant current, as is the case for the trough of the alternating current and cathodal constant current. The effects of alternating currents are similar to those of the analogous constant current paradigm, indicating that plasticity modulation is consistent with the instantaneous incremental membrane polarization on a millisecond timescale. LTP induction is applied at the 20 minute mark. All data are normalized to the mean of the 20 baseline responses before induction and are represented as mean±s.e.m across slices.

### Electric field interacts with plasticity induction on millisecond timescale

Membrane polarization during DCS has been well characterized (17–22) and is well described by cable models of stimulated neurons (Figure 1A)(21–23). We previously argued that the effects of DCS on tetanus-induced LTP are due to membrane polarization (14). If this is the case for TBS-induced LTP as well, then there is no need for the DCS to be constant over long periods of time. It would suffice for the DCS field to coincide with TBS synaptic inputs on the time scale of the neuronal membrane time constant (e.g. 30ms)(18). To test for this, we applied theta-frequency alternating current stimulation (sinusoidal 5 Hz at 20 V/m) during TBS induction. The peak phase of this alternating current corresponds to the same electric field as anodal DCS, while trough corresponds to cathodal DCS. When TBS bursts were timed to coincide with the peak of the alternating current, LTP was enhanced, as with anodal DCS (Figure 1C; control: 1.287+−0.025, N=52; peak: 1.467+−0.093, N=9, p=0.014; N here and below indicates the number of slices). TBS timed to the trough of the alternating current had no significant effect on LTP, as with cathodal DCS (Figure 1C; trough: 1.184+−0.035, N=6, p=0.173). These data suggest that the electric field need only coincide with potentiating synaptic input on the millisecond timescale, and does not require any prolonged buildup of DCS effects in order to affect LTP. This is consistent with the notion that instantaneous membrane polarization due to DCS is what interacts with synaptic activity to modulate the resulting plasticity (14).

### Effect of DCS on LTP is pathway specific

Hebbian synaptic plasticity is classically characterized as a pathway specific process, i.e. only pathways that are coactive with the postsynaptic neuron are strengthened (35). Our proposal that DCS enhances LTP through membrane potential implies that the effects of DCS should follow this pathway specificity. We tested this by monitoring two independent synaptic pathways in CA1 (Figure 2A). During induction, the strong pathway received TBS while the other inactive pathway was not stimulated. As expected, LTP was observed in the strong pathway (Figure 2B black; 1.377+−0.052, N=16, p=2.8E-6), but not the inactive pathway (Figure 2B gray; 0.986+−0.031, N=14, p=0.657), demonstrating the well-established pathway specificity of LTP (35). When this induction protocol was paired with anodal DCS, LTP was enhanced only in the strong pathway (Figure 2B red; 1.613+−0.071, N=14, p=0.011 vs. control), while the inactive pathway was unaffected (Figure 2B light red; 0.971+−0.028, N=14, p=0.724 vs. control), showing that the effects of DCS is specific to the potentiated pathway.

**Figure 2.**
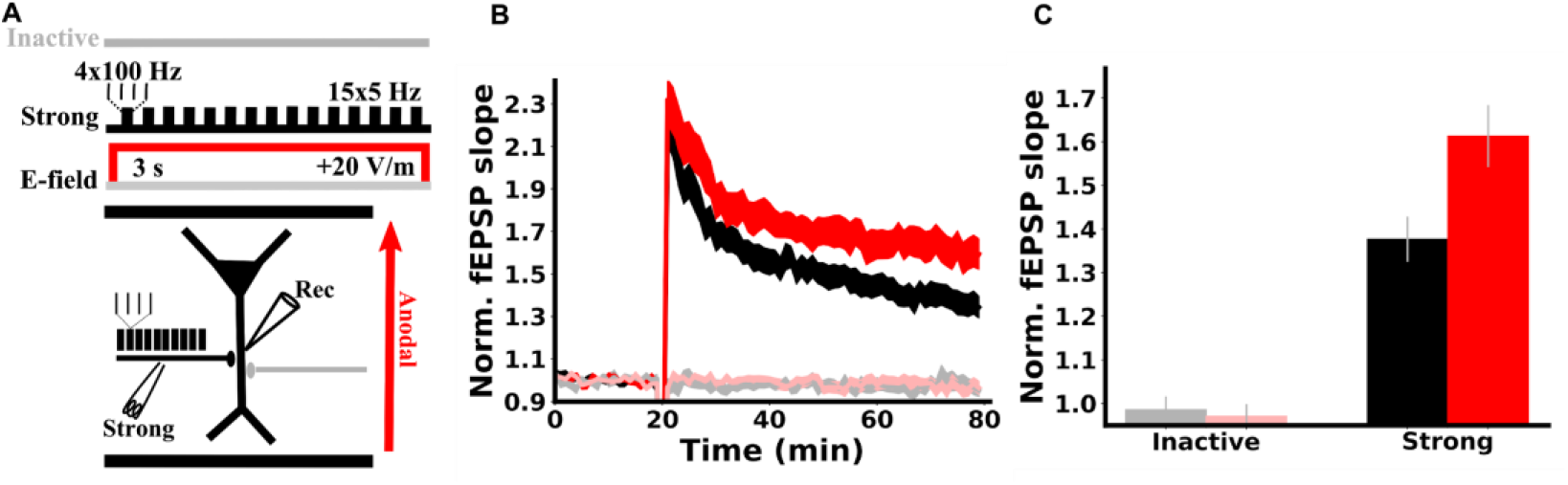
DCS effect is specific to the potentiated pathway. **A)** Schematic of the experimental setup. Two synaptic pathways are monitored before and after plasticity induction. During induction, one pathway is activated with TBS (black, strong), while the other pathway is inactive (grey), and anodal DCS is applied across the slice throughout the duration of induction (3 s, red). **B)** Plasticity is pathway specific and so are DCS effects. LTP was observed only in the pathway that received TBS (black trace), demonstrating pathway specificity. Anodal DCS enhanced LTP only in the potentiated pathway (red vs black) and had no effect on the inactive pathway (light red vs. gray), upholding Hebbian specificity. fEPSP slopes are normalized to the mean of the 20 of baseline responses prior to induction. Induction is applied at the 20 minute mark. **C)** Summary of pathway specific effects of DCS. The mean of the last 10 normalized slopes (51-60 min after induction) are used for each slice. Data are represented as mean±s.e.m across slices.

### DCS boosts Hebbian associativity

Another important property of Hebbian plasticity is pathway associativity, which is a cellular mechanism thought to underlie the formation of cell assemblies and associative learning (35–37). Pathway associativity refers to the potentiation of separate synaptic pathways arriving onto the same postsynaptic neuron when they cooperate to drive the postsynaptic cell. For example, a synaptic input that is too weak on its own to induce plasticity can undergo plasticity if it is coactivated with a strong input that helps to drive the postsynaptic cell.

We tested how DCS affects Hebbian associativity by again monitoring two synaptic pathways. First, only a weak input (15 pulses at 5 Hz) was used during induction (Figure 3A). In the absence of DCS, no lasting plasticity was observed in this weakly activated pathway (Figure 3A gray; 0.998+−0.041, N=13, p=0.966) or the other inactive pathway (Figure 3A black; 0.958+−0.037, N=13, p=0.275). DCS also had no effect on the weak (Figure 3A light red; 1.041+−0.038, N=13, p=0.445) or inactive pathway (Figure 3A red; 0.963+−0.011, N=13, p=0.908). This result further confirms the specificity of DCS effects, in that pathways that are not undergoing plasticity are unaffected by DCS.

**Figure 3.**
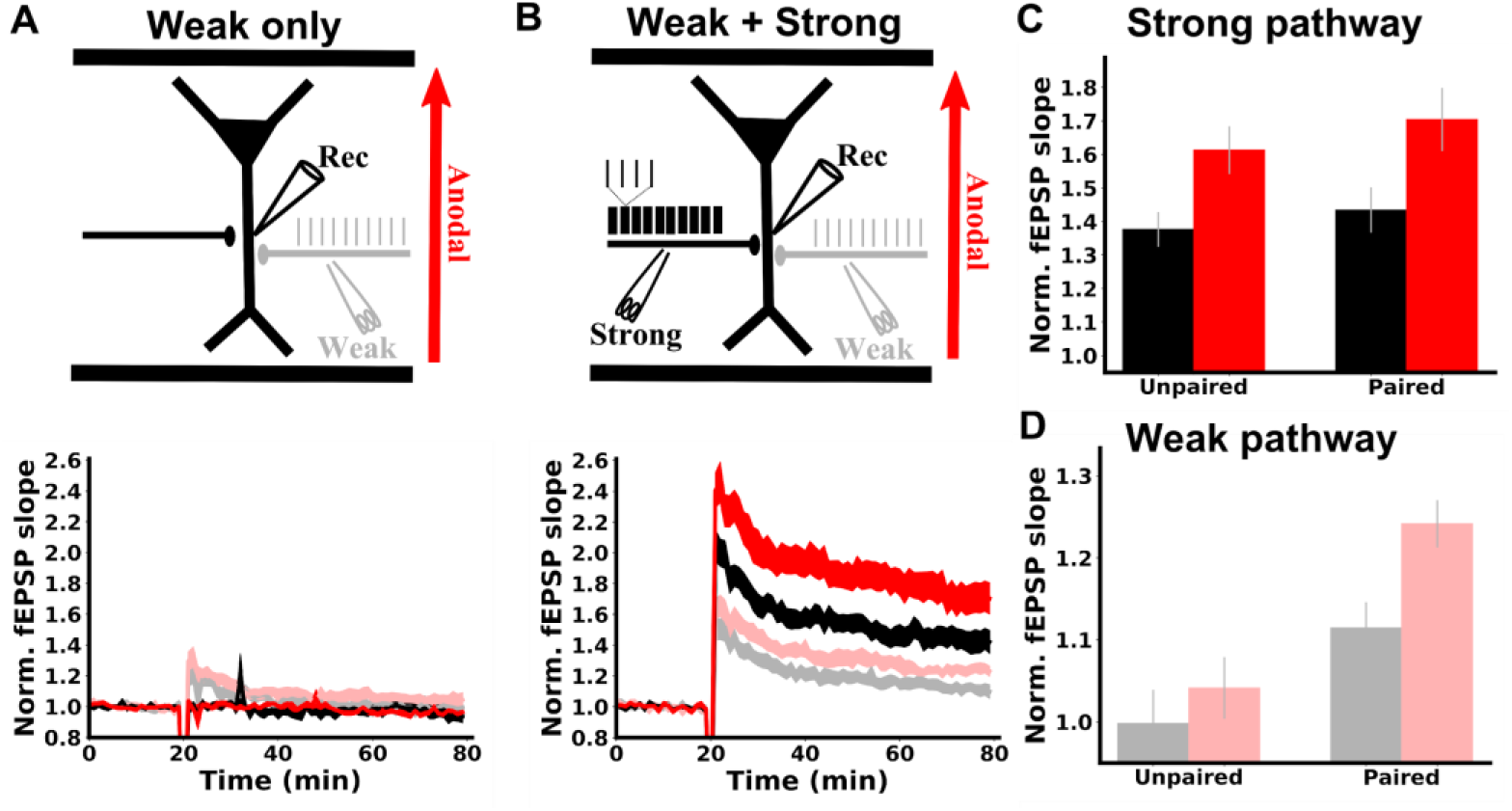
DCS enhances associativity between synaptic pathways. **A) Top**: schematic of experimental design. Two synaptic pathways were monitored. During induction, one pathway was weakly activated at 5 Hz with 15 pulses (grey), while the other pathway was inactive (black). Anodal DCS was applied throughout induction (3 s, red). **Bottom:** weak synaptic activation had no lasting effect on synaptic strength in either pathway with DCS (red, light red) or without DCS (grey, black). **B) Top:** schematic of experimental design. Again, two synaptic pathways were monitored. Now during induction, one pathway was activated with a TBS protocol (strong, black). The other pathway was activated with 15 pulses at 5 Hz (weak, grey). Weak pathway pulses were temporally aligned to the second pulse in each TBS burst. **Bottom:** without DCS, the strong pathway was potentiated (black) and the weak pathway was now also potentiated (grey), demonstrating associative plasticity between these pathways. With DCS, LTP was enhanced in the strong pathway (red) and the weak pathway (light red), demonstrating that the associativity between pathways was enhanced. **C)** Summary of LTP experiments in the strong pathway. Pairing with the weak pathway did not increase strong pathway LTP, and DCS had a similar effect on LTP in both cases. **D)** Summary of LTP experiments in the weak pathway. fEPSP slopes are normalized to the mean of the 20 of baseline responses prior to induction. Induction is applied at the 20 minute mark in panels **A,B**. The mean of the last 10 normalized slopes (51-60 min after induction) are used for each slice in panels **C,D**. Data are represented as mean±s.e.m. across slices.

In a second experiment, the weak input is now paired with a strong input (TBS) during induction (Figure 3B). During induction, weak pathway inputs are timed to arrive at precisely the same time as the second pulse of each theta burst. This induces LTP in the strong pathway as before (Figure 3B black; 1.435+−0.067, N=13, p=3.1E-5), but now the weak pathway is also potentiated (Figure 3B gray; 1.115+−0.031, N=13, p=0.003), replicating classic associativity between the two pathways (35). If this protocol is paired with DCS during induction, LTP is now boosted in both the strong (Figure 3B red c.f. black; 1.705+−0.094, N=13, p=0.029) and the weak pathway (Figure 3B light red c.f. gray; 1.242+−0.029, N=13, p=0.006). DCS therefore enhances the Hebbian associativity between the strong and weak pathways (Figure 3D). We note that plasticity was similar in the strong (TBS) pathway, regardless of whether it was paired with the weak pathway (Figure 3C black), and that the effect of DCS on the strong pathway was indifferent to pairing as well (Figure 3C red).

### Effects are consistent with DCS modulation of somatic spiking

We hypothesized that the effects of DCS on TBS-induced LTP are due to membrane polarization. However, DCS will alter membrane potential in both the soma and apical dendrites of pyramidal neurons, but with opposite polarities (18,23). We therefore aimed to test whether the effects of DCS on LTP were consistent with somatic or dendritic membrane polarization. To do so, we took a similar approach as in previous work (14). LTP was induced by stimulation of Schaffer collaterals with TBS in either apical or basal dendritic compartments of CA1 (Figure 4B). DCS is expected to have opposite effects on dendritic membrane potential in basal as opposed to apical dendrites (Figure 5A)(18,23). Effects due to DCS-induced *dendritic* polarization should therefore be *opposite* when synapses are activated in apical or basal dendrites. However, effects due to DCS-induced *somatic* polarization should be the *same*, regardless of the location of synaptic activation (i.e. there is only one soma per neuron). Therefore, observing different effects in apical and basal compartments would rule out somatic polarization as a main determinant of the plasticity modulation.

**Figure 4.**
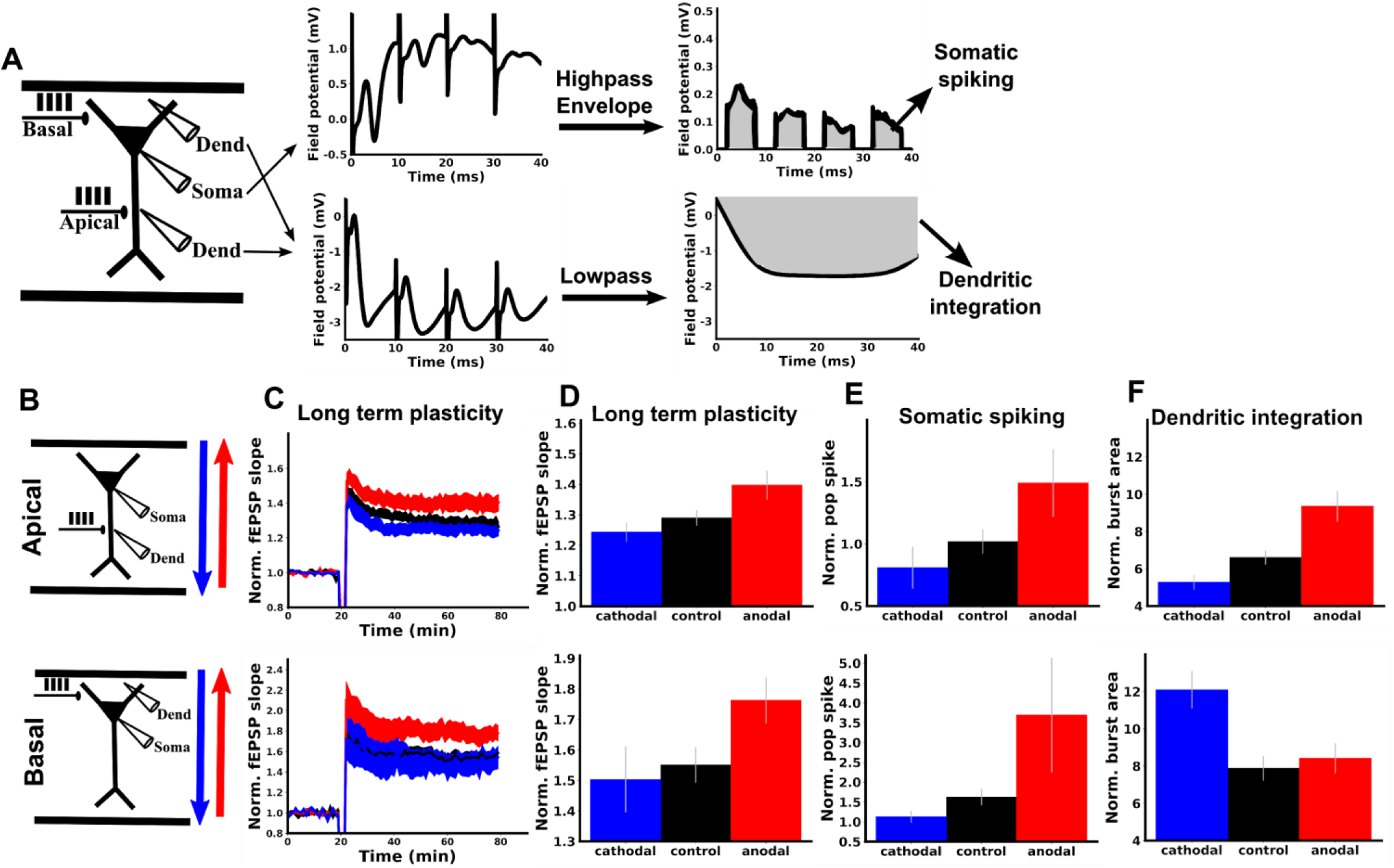
DCS modulation of TBS-LTP is consistent with modulation of somatic spiking rather than dendritic integration. LTP was induced with TBS in either apical (**top row, B-F)** or basal (**bottom row, B-F**) dendritic regions of CA1. TBS induction was paired with anodal (red), cathodal (blue), or no DCS (black). **A**) Schematic of experiments and methods for deriving somatic and dendritic activity metrics. For both apical and basal protocols, one recording electrode was placed in the dendrites (Dend) near the bipolar stimulating electrode (Apical or Basal) and one electrode was placed near the CA1 somatic layer (Soma). Examples of raw voltage traces from each recording electrode during a single burst of the induction protocol are displayed in the middle panel. To derive a measure of dendritic integration, the dendritic recording was low-pass filtered, and the integral of this filtered signal was taken for each burst during TBS (gray area). To derive a measure of somatic population spiking, the somatic recording was high-pass filtered, and the integral of this signal’s envelope during each burst was used (gray area; excludes periods of stimulation artefacts; see methods). **B)** Schematic of apical (top row) and basal (bottom row) experiments. **C)** Anodal DCS (red) boosts LTP in both and apical and basal dendrites compared to control (black). Cathodal DCS (blue) had no significant effect in either apical of basal dendrites. TBS was applied with or without DCS at the 20 minute mark. Note that the top panel is identical to Figure 1A (shown again here for comparison). **D)** Summary of the data in C. The mean of the last ten normalized responses were used for each slice. **E)** Population spiking measured for the first bipolar input pulse of the last burst (see Supplemental Figure S2C for all pulses during induction). **F)** Population dendritic integration for the last burst of TBS (see Supplemental Figure S2F for all bursts during induction). All data normalized to the mean of the 20 baseline responses before induction and error bars represent standard error of the mean.

**Figure 5.**
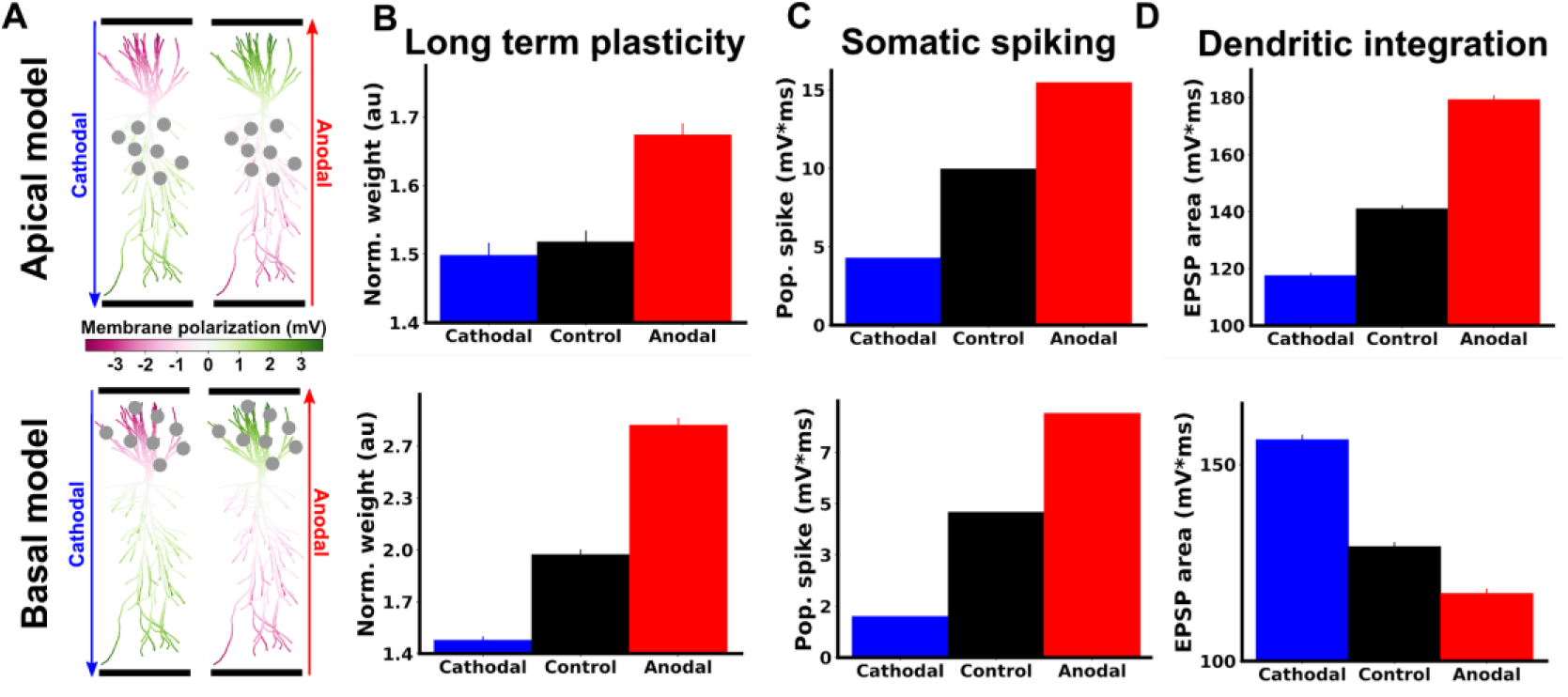
Model captures the effects of DCS on long term potentiation, somatic spiking and dendritic integration. **A)** Membrane polarization throughout model pyramidal neuron in response to 20 V/m anodal (red) or cathodal (blue) DCS. Green compartments are depolarized due to DCS, while magenta compartments are hyperpolarized by DCS. Gray circles indicate the location of synapses in apical (top row) or basal (bottom row) compartments that are activated with TBS. **B)** Model predictions of changes in synaptic weights qualitatively match LTP experiments (c.f. Figure 4D). The vertical axis (Norm. weight) is the average weight of all activated synapses at the end of simulation, calculated offline using the learning rule (41). **C)** Effects of DCS on somatic activity qualitatively match experimental measurements (c.f. Figure 4E). The vertical axis is the average across all neuron somas of the integral of the high-pass filtered voltage envelope (see methods). **D)** Effects of DCS on dendritic integration qualitatively match experimental measurements (c.f. Figure 4F). The vertical axis is the average across all recorded dendritic locations of the high-pass filtered envelope of the voltage (see methods).

Here we found that DCS had the same effect on LTP in both basal and apical dendrites (Figure 4C,D). This result is consistent with plasticity effects of DCS being driven primarily by effects on somatic spiking. To further test this, we looked at measures of dendritic integration and somatic spiking in each condition (Figure 4A, see Methods for details of analysis). Indeed, we found that DCS had a similar effect on somatic spiking (Figure 4E), but opposite effects on dendritic integration in apical versus basal dendrites (Figure 4F). Thus, the effect of DCS polarity on LTP mirrors that of the effect on the soma, but not dendrites.

### Computational model

To further understand how changes in membrane potential due to DCS lead to the observed changes in plasticity, we turned to a computational model. We modeled a CA1 pyramidal neuron based on a previously validated biophysical model, using the NEURON software package (38–40). To simulate the effects of DCS, we applied a uniform extracellular electric field (voltage gradient) with NEURON’s extracellular mechanism (23). This extracellular field is known to polarize the cellular membrane with opposite polarities in apical and basal compartment (Figure 5A)(18). To calculate activity-dependent synaptic plasticity, we used a voltage-based plasticity rule (41) that has been used previously to replicate a wealth of synaptic plasticity data (41–43). Here we manually selected parameters for this plasticity rule such that we could qualitatively reproduce canonical spike-timing dependent plasticity (STDP) experiments (26,43)(Supplemental Figure S1) and the effects of DCS on synaptic plasticity in our own TBS experiments (compare experiments of Figure 4D-F with model results of Figure 5). The model also reproduces the experimental results with alternating current stimulation (compare experiment of Figure 1C with model results of Figure S3). All simulation results that follow use the same parameters unless specified otherwise (Supplemental Tables).

### Associativity is enhanced through somatic spiking in simulations

Using the computational model, we then aimed to understand how DCS modulates TBS-induced LTP, while preserving specificity and associativity. Pathway specificity is explicitly built into the voltage-based plasticity rule of the model (41), following well established experimental results (35), namely synaptic weights are only allowed to change at active presynaptic inputs (see Methods). Since DCS does not by itself cause presynaptic activity, it cannot affect synaptic efficacy of the inactive pathway. Thus, the incremental membrane polarization due to DCS upholds Hebbian synapse specificity.

It is less clear however, exactly how DCS is able to boost associativity between the weak and strong pathways (Figure 3). We hypothesized that DCS boosted associativity through a boost of somatic spikes, which propagate to both weak and strong pathway synapses. To test this in the model, we simulated the experiments of Figure 3, by activating one pathway with TBS (strong) and the other pathway with the 5 Hz stimulation (weak). When the weak pathway was activated alone no spikes were generated and only very weak plasticity was observed (Figure 6D, weak only, black). Applying DCS in this case led to only minor changes in plasticity, as in our experiments (Figure 6D, weak only, compare red and black). However, when the weak input was paired with the strong input, action potentials were generated in the soma that back-propagated to weak pathway synapses (Figure 6B, black), and LTP was observed (Figure 6D, weak+strong, black). Therefore, the weak and strong pathway become associated by cooperating to produce somatic spikes, which are then shared by both pathways.

**Figure 6.**
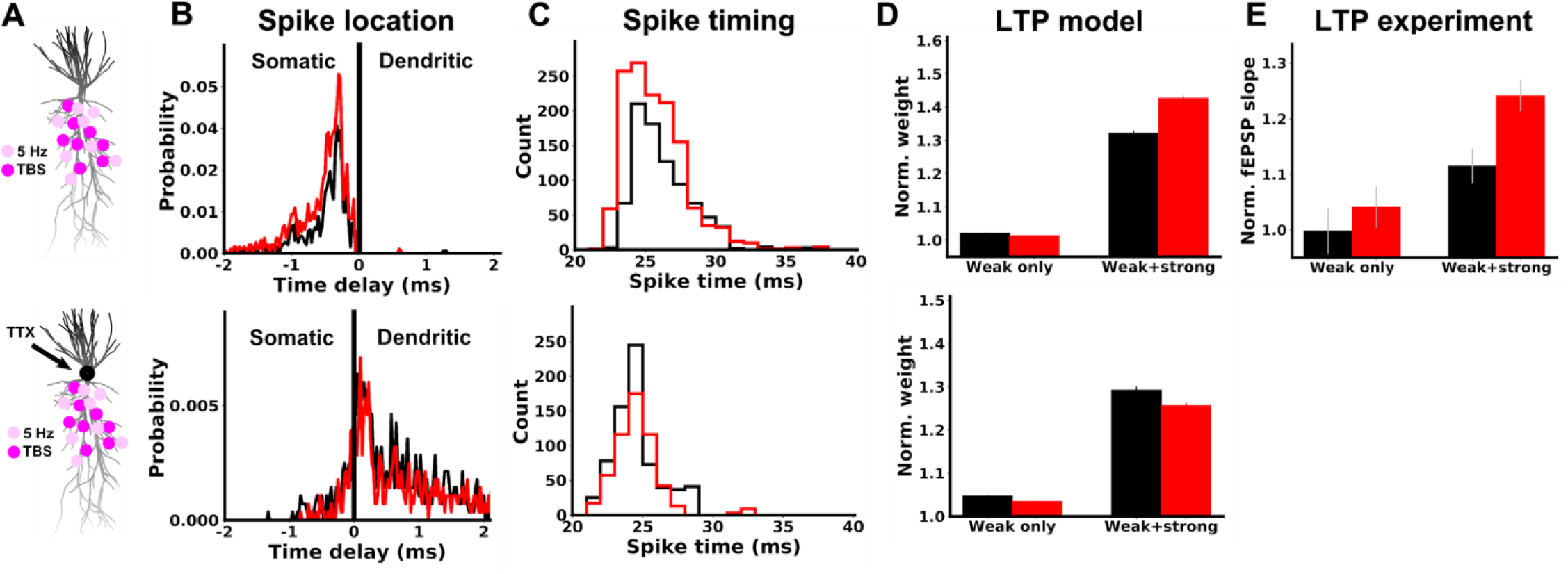
Boost of associative LTP is also explained by the effect of DCS on somatic spikes in computational model. **Top row: A)** Simulated neuron morphology, showing an example of how synapses are distributed in the weak (5 Hz, light pink) and strong (TBS, magenta) pathways. **B)** Distribution of time delays between spikes observed in the soma and at weak pathway synapses for 20 V/m anodal stimulation (red) or control (black). Negative time delays correspond to spikes that occur in the soma first. Due to variable propagation delays between synapses, it is possible for a spike initiated in the dendrite to reach the soma before other synapses. This produces a negative delay between the soma and these delayed synapses, even though the spike was dendritically initiated. It is not possible however, for a spike initiated in the soma to show a positive delay. **C)** Distribution of spike times recorded at all weak pathway synapses. Spike times are shown relative to the onset of the corresponding burst. **D)** Model prediction comparing plasticity in the weak pathway when it is unpaired (weak only) and paired (weak+strong). The vertical axis (Norm. weight) is the average weight of all weak pathway synapses at the end of simulation, calculated offline using the voltage-based learning rule (41). **E)** Experimental data (same as Figure 3D) shown again for comparison with panel D here. Both model and experiment show that anodal DCS increases LTP in the weak pathway only when it is paired with strong pathway activation. **Bottom row:** simulations and methods are identical to the top row, with two exceptions. First, we emulated the application of locally applied somatic TTX by setting voltage-gated sodium conductance to zero in the soma and axon, preventing the initiation of spikes in these compartments. Second, the number of synapses in each pathway was doubled, increasing the likelihood of spike generation, which now occurred in the dendrite. The testable prediction of the model is that in the presence of TTX now DCS will no longer boost LTP.

When strong and weak pathways were paired, DCS facilitated the initiation of somatic spikes (Figure 6B) and advanced their timing relative to the presynaptic input (Figure 6C), due to increased depolarization of the soma. This led to a boost in weak pathway plasticity only when paired with the strong input (Figure 6D, weak+strong), as observed experimentally (Figure 6E; same as Figure 3D).

To further validate the role of somatically initiated spikes in generating this DCS effect, we repeated the previous simulations, but set the voltage-gated sodium conductance to zero in the soma and axon (Figure 6A bottom). This is analogous to the local application of TTX at the soma (44), preventing the initiation of spikes there. If the strength of synaptic stimulation is increased, spikes can still be generated, but they initiate locally in the dendrite (Figure 6B bottom). Anodal DCS now reduces the probability of these spikes (Figure 6B bottom) and delayed their timing relative to the weak pathway input (Figure 6C bottom), due to DCS-induced hyperpolarization of the apical dendrites. A prediction of this model is therefore that TTX applied locally at the soma, would cause anodal DCS to have the opposite effect on pathway associativity (Figure 6D bottom), namely anodal DCS weakens rather than boosts LTP.

Taken together, the results of Figure 6 suggest that DCS can enhance associativity by facilitating the initiation of somatic spikes. The additional spikes can spread to synapses in both pathways and increase LTP, leading to a stronger association between the pathways.

### Interaction between synapse location and induction protocol

In a previous study, we used 20 Hz tetanic stimulation to induce LTP. We observed that a boost in LTP required opposite DCS polarities for apical and basal dendrites, suggesting that dendritic rather than somatic effects were dominant for this protocol (14), (Figure 7B, top two rows). This appears inconsistent with the previous claim that DCS effects are mediated primarily through somatic spiking (Figures 4–6). However, the computational model can readily reconcile these results if we consider the different endogenous membrane voltage dynamics during 20 Hz tetanus and TBS protocols. For 20 Hz tetanic stimulation, inputs arriving early in the tetanus may elicit somatic spiking (Figure 7D), but these inputs quickly become subthreshold due to short term synaptic depression (Figure 7E). Since the majority of input pulses remain subthreshold, plasticity at these synapses is dominated by the local subthreshold dendritic potential (Figure 7F). Because the DCS-induced polarization is opposite in apical and basal dendritic compartments, the effects on plasticity are also opposite there (Figure 7B,C; compare 20 Hz apical to 20 Hz basal).

**Figure 7.**
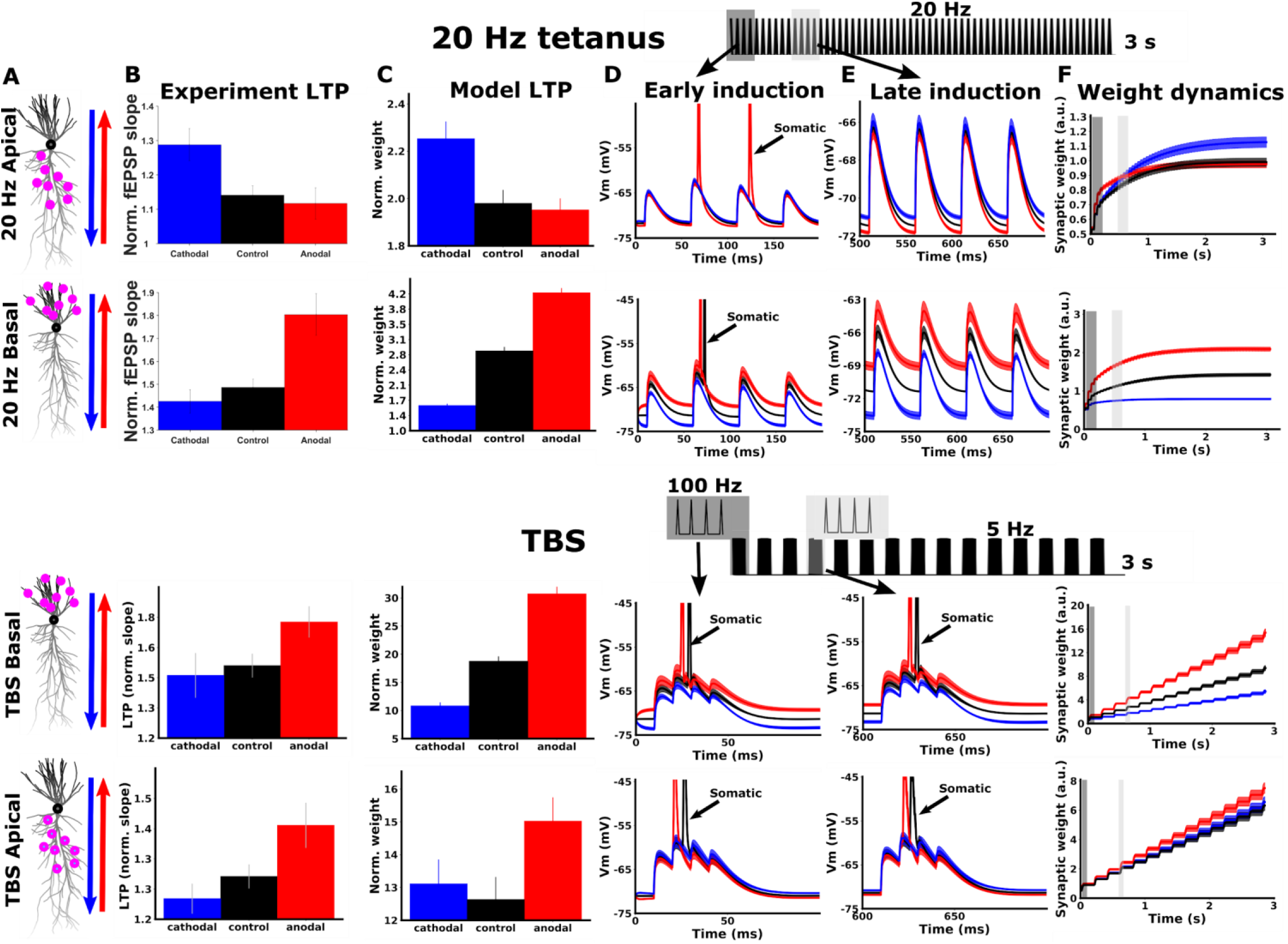
Model captures interaction between dendritic location and induction protocol. **A)** Simulated neuron morphology, showing distribution of activated synapses for 20 Hz (top two rows) and TBS (bottom two rows). Arrows indicate the direction of the DC fields for anodal (red) and cathodal (blue) stimulation. **B)** Experimental LTP results for each condition. The vertical axis is the average of the last ten normalized fEPSP responses. The top two panels are reproduced from data in (14). The bottom two panels are identical to figure 4D, shown again here for comparison. **C)** Model LTP predictions qualitatively match experimental LTP results (c.f. **C**; same direction of DCS effect). The vertical axis (Norm. weight) is the average weight of all activated synapses at the end of simulation, calculated offline using the learning rule of (41). **D)** Example simulated voltage traces for individual cells recorded only at activated synapses during the first four input pulses. Traces are averaged over all activated synapses for the example cell. Spikes that back-propagate from the soma are indicated with arrows. **E)** Same as D, but at a later time point in the simulation (pulses 10-13 for 20 Hz tetanic stimulation; pulses 13-16 for TBS simulations). Note that for 20 Hz stimulation synaptic depolarization is reduced due to short term depression and somatic spiking ceases very early in the simulation. During this subthreshold period, DCS causes a small shift in membrane potential and the resulting plasticity. Since DCS causes opposite subthreshold polarization in apical and basal dendrites, the effect on LTP is also opposite in apical and basal dendrites (**C**, top two rows). For TBS simulations, recovery from short term depression between bursts allows bursts later in the simulation to produce somatic spikes. Plasticity throughout the simulation is controlled by somatic spikes, and is similar in apical and basal dendrites (**C**, bottom two rows) **F)** Dynamics of synaptic weights during the full simulation, averaged over the entire population of activated synapses. For TBS simulations, the weight change is approximately linear in the number of bursts, as each successive burst is equally effective at inducing plasticity. For 20 Hz stimulation, the weight change saturates with the number of pulses, as each successive pulse is weaker due to short term depression. Only the weight at the end of the simulation is used to predict the resulting LTP in experiments (**C**). Gray boxes in **F** indicate time periods for early (dark gray) and late induction (light gray) that are plotted in **D** and **E**, respectively. A schematic of the input pulse train and relative timing of early (dark gray) and late (light gray) induction pulses are shown at the top. All data are represented as mean±s.e.m.

During TBS on the other hand, each burst in the induction is close to threshold at the soma. Somatic action potentials are generated throughout the induction, and plasticity at each synapse is dominated by the back-propagation of these spikes (Figure 7D,E; bottom two rows). Effects of DCS on plasticity therefore follow the effects on somatic spike generation, regardless of the dendritic location of the synapses (Figure 7B,C; compare TBS apical to TBS basal). Indeed, our experimental recordings of somatic spikes and dendritic integration in the CA1 population support this notion (Figure 4). Performing a similar analysis in the model recapitulates this result (Figure 5, c.f. Figure 4D-F).

The results of Figure 7 highlight the complex interaction between endogenous synaptic input dynamics, synapse location, and DCS-induced polarization. Despite the complexity, Figure 7 also points to a simple and more general principle: when endogenous plasticity is primarily driven by *somatic* sources of depolarization (e.g. backpropagating somatic spikes), DCS-induced polarization at the *soma* determines effects on plasticity. This is what we observe with TBS (Figure 7 bottom two rows). When endogenous plasticity is primarily driven by *dendritic* sources of depolarization (e.g. subthreshold depolarization or dendritic spikes), DCS-induced polarization at the *dendrite* determines effects on plasticity. This is what we observe with 20 Hz tetanus (Figure 7 top two rows) or when we block somatic spiking (Figure 6 bottom row).

### Dose response and distribution of plasticity effects

We are ultimately interested in understanding the effects of weaker electric fields that occur in the human brain during clinical tDCS, which are on the order of 1 V/m (45,46). The model presented above is able to reproduce several experimental effects of DCS (Figures 5–7, Supplemental Figure S3) and canonical synaptic plasticity results (Supplemental Figure S1) with the same set of parameters (Supplemental Table 1). Because the model includes the actual morphology of CA1 pyramidal neurons, the electric field magnitude in simulations has a precise mapping to the electric field in experiments. We therefore used the model to make predictions for how weaker electric fields would influence synaptic plasticity.

We first measured the passive membrane polarization throughout the model neuron in response to DCS (Figure 8A). As observed experimentally (18), we found that the subthreshold membrane polarization is linear in the electric field magnitude, with opposite polarization in the soma and apical dendrites (Figure 8B). Next we repeated simulations of TBS with DCS at varying electric field magnitudes (+− 1, 2, 5,10,15,20 V/m). For a given electric field magnitude, we quantified the effect of DCS at each synapse and averaged over all active synapses in the apical dendrite (gray circles in Figure 8A; see Methods). We found that the mean effect of DCS on plasticity is monotonic in the electric field magnitude (Figure 8E). While each polarity of electric field produces an approximately linear dose response, we observed a greater slope for anodal (positive) electric fields. This asymmetry of anodal and cathodal effects is consistent with our experimental observations (Figure 1B).

**Figure 8.**
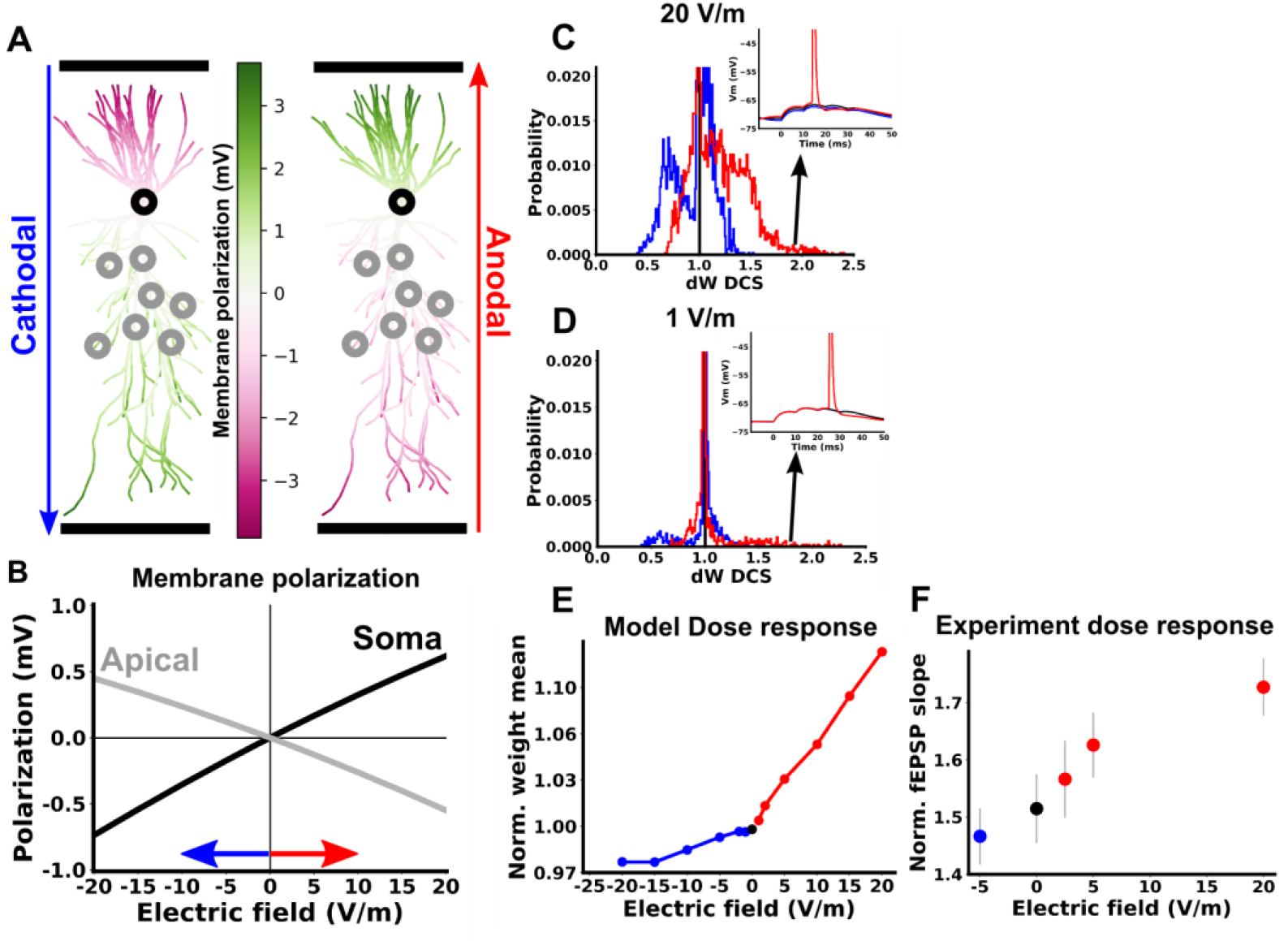
Dose response for computational model of TBS in apical dendrites. **A)** Membrane polarization of a CA1 pyramidal cell in response to 20 V/m cathodal (left) and anodal (right) electric field. **B)** Membrane polarization in response to varying electric field magnitude. On the horizontal axis positive values correspond to anodal DCS and negative values correspond to cathodal DCS. The gray curve is averaged over all segments in the apical dendrite, and the black curve is measured at the soma. **C,D)** Distribution of DCS effects on synaptic weight in response to TBS in apical dendrites. The horizontal axis is the the final synaptic weight during a simulation with DCS divided by the final synaptic weight in the same cell under control conditions. ΔW_DCS_ therefore measures the change in weight caused by DCS for each synapse. Inset shows example voltage traces for synapses in the tail of the distribution. These synapses correspond to cases where the control simulation brought the cell to slightly below threshold, such that DCS was able to cause firing and produce a large change in the weight. **E)** Mean of the synaptic weight change (ΔW_DCS_) due to TBS, averaged over all simulated apical synapses, as a function of DCS electric field. **F)** Experimental LTP as a function of DCS electric field. All data are represented as mean±s.e.m.

If we consider the distribution of DCS effects over all apical synapses, we find that for weak fields the mean effect of DCS is predominantly driven by the tail of this distribution, where very few synapses have large changes in plasticity (Figure 8D). For a small number of cells that are close to threshold, a weak field may cause a spike that would have otherwise not happened. This causes a large jump in all synaptic weights for a few highly sensitive cells. While most synapses see very small effects on their weights due to small effects on spike timing and subthreshold polarization, a small number of synapses experience a large effect on their weights due to the initiation of new spikes.

We next experimentally tested the dose response by varying the DCS electric field (−5, 0, 2.5, 5, 20 V/m; cathodal negative; anodal positive). Consistent with the prediction of the model, there was a monotonic relationship between electric field and the magnitude of LTP (Figure 8F): −5 V/m (1.41+−0.049, N=16), 0 V/m (1.46+−0.060, N=20), 2.5 V/m (1.52+−0.068, N=15), 5 V/m (1.57+−0.057, N=14), 20 V/m (1.67+−0.051, N=18), with larger effects for anodal stimulation (see also Figure 1B). Unlike the model however, the effect of anodal stimulation appears to be saturating at 20 V/m, perhaps reflecting saturation of LTP itself. This discrepancy can be accounted for by considering that synaptic weights in the model can grow without bound, unlike biological synapses (47).

## Discussion

Synaptic plasticity is critical for many forms of learning and tDCS has been thought to alter synaptic plasticity (6,48). How stimulation may interact with ongoing synaptic activity to alter plasticity remains poorly understood. Here we found that weak electrical stimulation with constant direct currents can enhance LTP, while maintaining input specificity and associativity. We propose a model in which DCS boosts endogenous Hebbian synaptic plasticity through modulation of pyramidal neuron membrane potential dynamics. As this model predicts, the effects of DCS also reflect the input specificity and input associativity of the endogenous Hebbian plasticity.

This framework produces a number of testable predictions for clinical experimentation. First, the efficacy of clinical protocols should improve when tDCS is paired with a learning task which induces plasticity, instead of the common practice of pairing tDCS with “rest”. Second, when tDCS is paired with a learning task, we postulate that the effects should be highly specific to the trained task. Finally, the pairing of tDCS with Hebbian plasticity and learning can be thought of as a method for functional targeting, since tDCS should only affect synaptic pathways that are already undergoing plasticity due to the paired task. This may alleviate the prevailing concern that focal stimulation of a desired target in the brain is not possible with transcranial electrical stimulation.

### Hebbian plasticity

Hebb originally proposed that coincident pre and postsynaptic *firing* was required for enhanced synaptic efficacy (49). Over time the concept of Hebbian plasticity has come to incorporate forms of plasticity that depend on correlated pre and postsynaptic activity variables, regardless of the exact biophysical implementation (50). While we do not directly measure or manipulate postsynaptic firing here, TBS-induced LTP at CA1 Schaffer collaterals has been shown to be Hebbian in that it depends on pre and postsynaptic activity and exhibits classic Hebbian properties of input specificity and associativity (51). The synaptic plasticity rule in our model is similarly Hebbian in that plasticity depends on correlated pre and postsynaptic activity in the form of presynaptic spike arrival and postsynaptic membrane voltage (41).

### Input specificity

Input specificity is a property of Hebbian plasticity whereby only synaptic inputs that are coactive with the postsynaptic neuron, and presumably relevant for the current task, are strengthened (35). The computational significance of this specificity has been recognized for some time, as it allows a network of neurons to learn sparse, non-overlapping neural representations (52). In practice, this is implemented in the brain by molecular machinery which responds to elevated activity specifically at task-relevant synapses (53). Here we show that DCS enhances LTP in a manner that respects this input specificity. DCS only boosts the strength of synapses that are active and already undergoing endogenous plasticity. Based on this observation, we make two predictions for the optimization of tDCS effects in humans.

First, tDCS effects in humans should similarly exhibit synaptic input specificity, which would be reflected as task specificity in the cognitive domain. Indeed, there is good evidence for task-specific effects of tDCS, despite its lack of spatial focality in the brain (54,55). This property may be central to the broad application of tDCS. It implies that tDCS can be used flexibly in combination with many different tasks and with limited side effects, despite stimulation reaching large regions of the brain. Second, tDCS effects may be most pronounced when paired concurrently with training that induces plasticity. Again, there is evidence for this in the human literature (33,56). It may be possible to leverage these properties further by pairing stimulation with forms of learning that are known to rely heavily on Hebbian mechanisms (57–59).

### Associativity

Associativity refers to the potentiation of a weak synaptic input when it is paired with strong input at other synapses to the same neuron. In this sense the weak input becomes associated with the strong input. This can serve as a cellular mechanism to bind previously unrelated information as in classical conditioning (60), and to form cell assemblies for associative learning (35–37). Here we show that anodal DCS can further enhance this associativity, which may manifest as an increased probability of forming associations between stimuli during learning that involves Hebbian plasticity. We did not explore associativity under cathodal DCS as we saw no effect on LTP for the single pathway experiment, but we cannot rule out an associative effect of cathodal DCS.

### Asymmetry

As in our previous work (14,61) and in many tDCS studies (28,62,63), we observe asymmetric results with respect to DCS polarity. Anodal DCS enhanced LTP, while cathodal DCS had no discernible effect with the current sample sizes. This stands in contrast to the symmetric membrane polarization observed with opposing field polarities (18). Of course, the brain exhibits highly nonlinear responses to changes in membrane voltage, from the level of ion channels to the propagation of activity in a recurrent network. In this sense, it is perhaps not surprising that responses to DCS are nonlinear. However, it remains a crucial topic to understand which sources of nonlinearity are most relevant for DCS, and whether these persist in human tDCS. Below we speculate on some of these potential sources, although we are unable to disambiguate them here, as it is beyond the scope of the current study.

The asymmetry may result from the interaction between DCS effects on different neuronal compartments. For example, during cathodal stimulation, *depolarization* of *apical dendrites* can counteract *hyperpolarization* of *somas* so that there is no reduction in LTP (61)(Figure 8A,B). However, this mechanism cannot explain the asymmetry we observed for LTP in basal dendrites (Figure 4C,D; bottom row), as the direction of polarization is the same in both basal dendrites and somas (Supplemental Figure S5A,B). While our model does predict a nonlinear dose response in basal dendrites (Supplemental Figure S5E), this is more likely due to nonlinear responses of voltage-gated ion channels or the synaptic plasticity molecular machinery.

A nonlinear dose response may also result from the distribution of initial synaptic states in the cell population that we record from. For example, the prior history of the recorded synapses may be such that they are biased towards an increase in strength (64). Similarly, it could reflect the distribution of cell excitability, such that cells are biased toward an increase in firing. With this in mind, we analyzed the input-output relationship between fEPSP’s and population spikes in our baseline recordings. Indeed, we found that our experiments are run near a nonlinearity in this input-output relationship, such that population spiking could be more readily increased than decreased (Supplemental Figure S4).

Mirroring the asymmetric effect of DC polarity, the effects with respect to phase of AC stimulation was also asymmetric. This suggests that even in the absence of information about the precise timing of synaptic inputs when tACS is applied in humans, a net enhancement of LTP may be expected when tACS is paired with synaptic plasticity induction. Notably, the boost in LTP was also larger here for ACS than DCS, perhaps owing to the frequency response properties of pyramidal neuron membranes showing a peak at theta frequencies (20,65).

### Mechanism

Perhaps the most well characterized cellular effect of electrical stimulation is the modulation of somatic membrane potential and firing probability (18,20,61,65–70). In human tDCS studies, it is the modulation of motor-evoked potentials, which have been linked to long-term plasticity (48,71,72). Here we propose a model which translates acute changes in firing probability and timing into long term changes in synaptic plasticity. In addition to several other phenomena (7,8,73,74), previous studies have pointed to the effects of DCS on BDNF release (12,13,15,75). While the precise mechanisms remain unclear, BDNF appears to be released in response to postsynaptic depolarization and involved in LTP induction (76–78). BDNF may therefore be an essential part of the molecular machinery that translates DCS-induced effects on membrane potential dynamics into changes in plasticity.

Electric fields are also known to alter cell motility and immune responses (79,80). However, these effects unfold over the course over many minutes to hours. During prolonged stimulation, it is likely that various effects on cellular physiology begin to take hold simultaneously, with interactions between them. However, robust effects were generated here with remarkably short stimulation duration (3 s), which depended on stimulation polarity with sub-second timing (100 ms, Figure 1C). Polarization of neuronal membranes is the only known effect of stimulation that acts on these timescales, making it a likely source of effects here. Inhibitory neurons were not included in our model as the effects of DCS are expected to be small, at least for neurons with symmetric morphology (19). However, we cannot rule out that DCS polarizes inhibitory neurons on a rapid time scale, which in turn may affect plasticity either directly or through network effects. Prolonged stimulation necessarily includes effects operating on both short timescales (membrane polarization, plasticity induction) and longer timescales (cell motility and immune responses), e.g. (81). However, shortening the stimulation and pairing it with quicker (sub-minute) bouts of training as we have done here, could be a useful strategy to isolate effects based on Hebbian plasticity induction, which operate on faster timescales.

Our experiments and computer simulations support a model in which DCS affects TBS-induced LTP primarily by somatic polarization and changes in somatic spiking (Figures 4–6). However, DCS-induced dendritic polarization is also likely to contribute to plasticity, as we suspect for 20 Hz tetanus experiments (Figure 6)(14). Our computational model can reconcile these results by considering the voltage dynamics during induction (Figure 6).

We propose a general principle that emerges from this result: when endogenous plasticity is primarily driven by *somatic* sources of depolarization (e.g. spikes), DCS-induced polarization at the *soma* determines effects on plasticity. When endogenous plasticity is primarily driven by *dendritic* sources of depolarization (e.g. subthreshold depolarization or dendritic spikes), DCS-induced polarization at the *dendrite* determines effects on plasticity. The relative contribution of somatic and dendritic DCS effects, and therefore the overall effect on plasticity, is not always obvious. The spatial location and temporal pattern of active synapses (including background synaptic input), as well as neuromodulator concentrations and intrinsic excitability can all shift the endogenous voltage dynamics towards somatic or dendritic dominance. Computational models, such as the one presented here, can help in this regard by exploring how DCS interacts with this large parameter space of endogenous synaptic activity. This should be an important next step for future work.

### Brain region

While electric current does reach the hippocampus and subcortical structures during stimulation (82), tDCS is thought to primarily act on neocortex. Here we chose hippocampus as a model system for the wealth of studies on hippocampal synaptic plasticity and the much neater organization of input pathways. While not identical, many excitatory plasticity mechanisms are conserved in pyramidal neurons between cortex and hippocampus (83), making our observations here informative for cortex as well. Indeed, the plasticity rule used here in our model has also been used to describe plasticity at neocortical excitatory synapses (26,41,43). Of course, further work is needed to validate this relationship with respect to DCS effects. It is also worth noting that this work, in addition to other recent studies (12,15,84), motivates the hippocampus as a target for tDCS.

### Dose response

Here we used a 20 V/m electric field in order resolve effects with a reasonable number of animals. Electric fields in the brain during typical tDCS experiments are expected to be 1 V/m or less (45). While we do not measure effects with this intensity, our computational model predicts a monotonic relationship between the population-mean synaptic plasticity and electric field magnitude (Figure 8C). For a given DCS polarity, the model predicts a linear relationship between field magnitude and mean plasticity effects (Figure 8E). To first order this implies population mean effects of ~1% for fields of 1V/m (we observe ~20% effects for 20 V/m), in line with effect sizes observed for acute effects of DCS (70). However, experimentally we observe a saturation with increasing stimulation intensity (Figure 8F). This linear approximation may therefore underestimate effect sizes with weaker fields.

We also note recent efforts to increase stimulation intensity up to 6mA in humans by distributing current across multiple electrodes (85), which can achieve electric fields of 3 V/m in the brain (82). Given our estimates here, this would generate effects on synaptic plasticity of ~3%, notably affecting a few synapses most strongly (Figure 8D). Recent in vivo rodent work suggests that a motor learning task leads to potentiation of ~1-2% of synaptic spines in a given volume of cortex (86), which is comparable to what we expect tDCS to achieve. Effect sizes of tDCS on synaptic plasticity in humans are therefore likely to be in a behaviorally relevant range.

## Methods

All animal experiments were carried out in accordance with guidelines and protocols approved by the Institutional Animal Care and Use Committee (IACUC) at The City College of New York, CUNY (Protocol 846.3 and 2016-24).

### Brain slice preparation

Hippocampal brain slices were prepared from male Wistar rats aged 3–5 weeks old, which were deeply anaesthetized with ketamine (7.4 mg kg^−1^) and xylazine (0.7 mg kg^−1^) applied I.P., and sacrificed by cervical dislocation. The brain was quickly removed and immersed in chilled (2–6 °C) dissecting artificial cerebrospinal fluid (aCSF) solution containing (in mM): Choline chloride, 110; KCl, 3.2; NaH_2_PO_4_, 1.25; MgCl_2_, 7; CaCl_2_, 0.5; NaHCO_3_, 26; d-glucose, 10; sodium ascorbate, 2; sodium pyruvate, 3. Transverse slices (400 μm thick) were cut using a vibrating microtome (Campden Instruments) and transferred to a chamber containing a recovery aCSF at 34 °C: NaCl, 124; KCl, 3.2; NaH_2_PO_4_, 1.25; MgCl_2_, 1.3; CaCl, 2.5; NaHCO_3_, 26; d-glucose, 25; sodium ascorbate, 2; sodium pyruvate, 3. After 30 minutes in the recovery solution, slices were transferred to a holding chamber containing recording aCSF at 30 °C: NaCl, 124; KCl, 3.2; NaH_2_PO_4_, 1.25; MgCl_2_, 1.3; CaCl, 2.5; NaHCO_3_, 26; d-glucose, 25; for at least 30 minutes. Finally, slices were transferred to a fluid–gas interface chamber (Harvard Apparatus) perfused with warmed recording aCSF (30.0 ± 0.1 °C) at 2.0 ml min^−1^. Slices were allowed to acclimate to the recording chamber for at least 30 minutes before recording started. The humidified atmosphere over the slices was saturated with a mixture of 95% O_2_–5% CO_2_. All aCSF solutions were bubbled with a mixture of 95% O_2_–5% CO_2_. Recordings started approximately 2 h after the animal was sacrificed.

### fEPSP recordings

Field excitatory postsynaptic potentials (fEPSP’s) were evoked using a platinum–iridium bipolar stimulating electrode placed in either stratum radiatum (apical experiments) or stratum oriens (basal experiments) of CA1 within 200 μm of the somatic layer. Recording electrodes were made from glass micropipettes pulled by a Sutter Instruments P-97 and filled with recording aCSF (resistance 1–8 MΩ). A “dendritic” recording electrode was placed in stratum radiatum (apical) or stratum oriens (basal) approximately 400 μm from the stimulating electrode in CA1 to record fEPSP’s. The stimulating electrode and dendritic recording electrode were placed at approximately the same distance from the CA1 somatic layer. For all experiments, a second “somatic” recording electrode was placed in the CA1 somatic layer to record population spikes. For two-pathway experiments (Figures 2 and 3), a second stimulating electrode was placed on the opposite side of the recording electrode.

fEPSP’s were quantified by the average initial slope, taken during the first 0.5 ms after the onset of the fEPSP. The bipolar stimulus intensity was set to evoke fEPSP’s with 30-40% of the maximum slope, which was determined at the onset of recording. Baseline fEPSP’s were recorded once a minute for at least 20 minutes before any plasticity induction was applied and only if a stable baseline was observed. For two pathway experiments, stimulation of each pathway was interleaved with an offset of 30 s. After plasticity induction, fEPSP’s were again recorded once per minute for 60 minutes. To measure synaptic plasticity, all fEPSP slopes were normalized to the mean of the 20 fEPSP’s immediately preceding induction. The amount of LTP in each slice is quantified as the mean of the last 10 minutes of normalized responses (51-60 minutes after induction). Group data are reported as mean and standard error of the mean across slices. Number of slices is indicated with variable N wherever statistics are reported.

### DCS

Uniform extracellular electric fields (±20 V/m) were generated by passing constant current (D/A driven analog follower; A-M Systems, WA, USA) between two large Ag-AgCl wires (1 mm diameter, 12 mm length) positioned in the bath across the slice starting 0.5 s before the onset of TBS and ending 0.5 S after the end of TBS (4 s total). Slices were oriented such that the somato-dendritic axis of CA1 pyramidal neurons was parallel to the electric field between the DCS wires (Fig. 1A). We name each polarity of DCS based on the orientation of the field relative to CA1 pyramidal neurons, and how pyramidal neurons are expected to be polarized. Here, anodal DCS depolarizes CA1 pyramidal neuron somas as it is expected to do in cortical pyramidal neurons under an anode in tDCS. Cathodal stimulation refers to the opposite polarity. Before each recording, DCS current intensity was calibrated to produce a 20 V/m electric field across each slice (typically 100–200 μA) by adjusting the current so that two recording electrodes separated by 0.8 mm in the slice measured a voltage difference of 16 mV (16 mV/0.8 mm = 20 V/m).

### Quantifying population somatic activity

To quantify the amount of somatic activity in response to synaptic input we used the following method. Raw voltage data recorded in the somatic layer was filtered with a 300 Hz highpass ARMA filter. The filter was designed using the butterworth algorithm via the signal.iirdesign function in the scipy package (design parameters: fs=10000; nyquist=fs/2; wp=300/nyquist; ws=200/nyquist; gpass=1; gstop=20; ftype=’butter’). We then defined somatic activity for each evoked response, *s*, as the integral of the high frequency envelope:

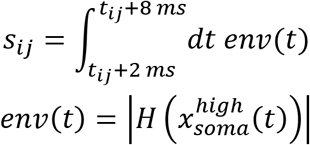

where 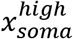 is the highpassed extracellular voltage recorded in the somatic layer, and *H*() is the hilbert transform calculated in python using signal.hilbert from the scipy package. *t*_*ij*_ is the onset time for the *i*^*th*^ input of the *j*^*th*^ burst, where *i* ∈ {1,2,3,4}and *j* ∈ {1,2, …15}. The somatic activity was calculated as the integral of this high frequency envelope in the time window 2-8 ms after *t*_*ij*_, chosen to avoid including the bipolar stimulus artifact. Somatic activity was then normalized to the mean of baseline values (20 responses prior to induction). The same method was used to calculate somatic activity in the population of model neurons (Figure 5C), except the recorded extracellular voltage in the somatic layer was replaced with the intracellular somatic voltage averaged over all simulated model neurons.

### Quantifying population spike timing

We expected that DCS would cause a shift in the average spike timing in the population during TBS (Supplemental Figure S2E). To create a measure of the mean spike timing, we performed a center of mass calculation on the somatic activity envelope

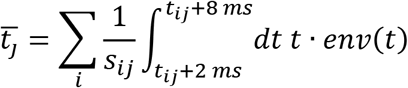

where 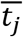 is the population spike timing of the *j*^*th*^ burst, *env* is the envelope of the highpassed extracellular voltage (see above), *t*_*ij*_ is the onset time for the *i*^*th*^ input of the *j*^*th*^ burst, where *i* ∈ {1,2,3,4} and *j* ∈ {1,2, …15}, and *s*_*ij*_ is the somatic activity as in the previous section. Again, we restrict the integrals to between 2 and 8 ms after each input pulse to avoid contributions of the bipolar stimulus artifact. 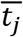 can be thought of as the temporal center of mass of the somatic activity during a burst. If more neurons in the population fire earlier during the *j*^*th*^ burst, then 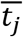 should decrease.

### Quantifying dendritic integration

To quantify the amount of dendritic integration in response to synaptic input we used the following method. Raw voltage data recorded in the dendrites (either stratum radiatum or stratum oriens) was filtered with a 5-50 Hz bandpass ARMA filter. The filter was designed using the butterworth algorithm via the signal.iirdesign function from the scipy package (parameters: fs=10000; nyquist=fs/2; wp=[5/nyquist, 50/nyquist]; ws=[0.1/nyquist, 100/nyquist]; gpass=1; gstop=20; ftype=’butter’). We then defined dendritic integration, *d*, for each burst during TBS as the integral of the band-passed signal:

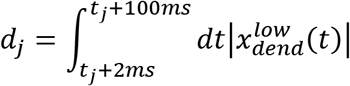

where 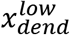 is the band-passed extracellular voltage recorded in the dendrite. For each evoked burst, *j*, the dendritic integration was calculated as the integral of this low frequency signal in the time window 2-100 ms after the onset of the burst, *t*_*j*_. Dendritic integration was then normalized to the mean of baseline values calculated for each fEPSP (20 responses prior to induction). The same method was used to calculate dendritic integration in the population of model neurons (Figure 5D), except the recorded dendritic extracellular voltage was replaced with the intracellular voltage averaged over all recorded dendritic segments in the simulated population of cells.

### Neuron model

Individual pyramidal cells were modeled in Python using the NEURON simulation package (87). To construct the model neuron, we reproduced the detailed biophysical neuron model of Migliore et al. (38), and then added parameter changes based on more recent studies. Unless otherwise specified, parameters are the same as in (38).

An L-type calcium channel was added throughout the cell as in (88). Sodium conductance in the axon was increased to replicate spike initiation in the axon initial segment (89). Synapses were set to have both AMPA and NMDA receptors, which were modeled as the difference of two exponentials. NMDAR conductance was modified by a voltage dependent mechanism as done previously (88,90,91). See supplemental tables for the full set of NEURON model parameters.

Synaptic conductances were modified by presynaptic short-term plasticity model as in (92). Specifically, AMPAR and NMDAR conductances were multiplied by a factor *A*, which captures short-term facilitation and depression dynamics at presynaptic terminals. *A* is the product of a facilitation variable *F*, and 3 depression variables *D*_*1*_, *D*_*2*_, *D*_*3*_

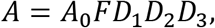

where A_0_ is a constant parameter, which we set to 1 at the start of simulations. At the time of each presynaptic spike, D is multiplied by a factor d such that

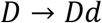

while a factor *f* is added to F such that

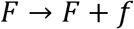

Both F and D decay exponentially back towards 1 between spikes according to

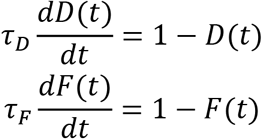

Each depression variable D_1_, D_2_, D_3_ follows the same dynamics, but with different parameters *τ*_*D*_ and *d*. The parameters of this short-term plasticity model were fit to the dynamics of fEPSP slopes during the various plasticity induction protocols in (14)(i.e. 0.5, 1, 5, 20 Hz trains) and during TBS in this study. The fit was constructed to minimize the squared error between values of *A* and the normalized fEPSP during induction using the `lsqcurvefit` function in matlab. See supplemental Table 2 for the resulting parameters.

The response of an individual pyramidal neuron to the bipolar stimulus in our brain slice experiments was modeled by randomly selecting a group of dendritic segments. AMPAR and NMDAR conductances were then activated simultaneously in the selected segments. In our experiments we expect that the bipolar stimulus will elicit this synaptic input in a population of pyramidal cells, with the number and location of synapses that are activated varying between cells.

For simplicity, we assume that an integer number of synapses ranging from 5 to 16 can be activated on each cell. This range was selected empirically so that the mean number of synapses produced somatic responses that were close to firing threshold during simulation of TBS. For each integer number of activated synapses, the synapses are randomly distributed on the dendrites, and this was repeated 20 times independently to create a population of 12*20=240 cells. For each cell, synapse locations were drawn randomly and with replacement from a uniform distribution over all dendritic segments that are allowed by the given experiment (e.g. basal dendrites or apical dendrites within 300 μm from the soma). By sampling with replacement, we allow multiple synapses to be activated on the same dendritic segment, mimicking the random activation of clustered synaptic inputs.

The electric field during DCS was modeled as uniform extracellular voltage gradient. The extracellular voltage at each point in space is then conveyed to each segment of the neuron by NEURON’s extracellular mechanism, as has been done previously (23). Since we are interested primarily in the effect of the extracellular field, for each simulation that applies an electric field there is a corresponding control simulation in which the NEURON model is identical except for the extracellular applied voltage. The effect of the applied field can therefore be compared to a precise counterfactual, where all other aspects of the model are identical.

### Voltage-based long-term plasticity rule

We are interested in how synaptic input and postsynaptic voltage dynamics during induction leads to long-term synaptic plasticity (and how DCS can modulate this plasticity). To simulate long-term synaptic plasticity in the model, we use the voltage-dependent plasticity rule of Clopath et al. (41). As done previously, we assume that actual changes in long-term synaptic strength are delayed relative to the induction period and do not contribute to the dynamics during induction. The plasticity rule is therefore used as a method to compute the final weight change expected at the end of induction and this calculation was done “offline”, after simulating the induction protocol. The synaptic weight change is calculated with the following rule (see (41) for further details), which requires information that is derived solely from presynaptic input arrival times and postsynaptic membrane potential measured locally at the synapse:

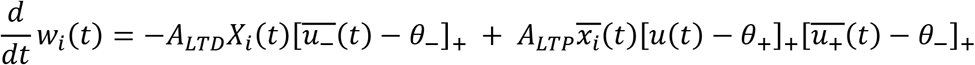

where *w*_*i*_ is the weight of the *i*^*th*^ synapse, *A*_*LTD*_ is a parameter that controls the rate of long-term depression (LTD), *A_LTD_* is a parameter that controls the rate of LTP, *X*_*i*_ is the presynaptic spike train, 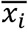 is a lowpass filtered version of the presynaptic spike train, *u* is the postsynaptic membrane potential measured locally at the synapse, 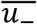 and 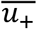 are lowpass filtered versions of the postsynaptic potential, *θ*_−_ is an LTD threshold parameter, *θ*_+_ is an LTP threshold parameter, and [·]_+_ indicates positive rectification. The dynamics of 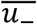, 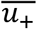, and 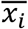 are given by

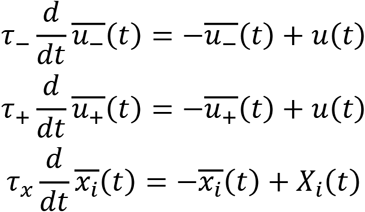

where *τ*_−_, *τ*_+_, and *τ*_*x*_ are time constants. We note that in the original learning rule of Clopath et al. (41), *ALTD* is a function of time and postsynaptic voltage, i.e. *A*_*LTD*_*(t,u)*, which implements homeostatic plasticity. Because we compute synaptic weight changes offline, a homeostatic mechanism is not needed to stabilize the voltage dynamics. To reduce the number of parameters of our model, we therefore treat ALTD as a constant. We apply a lower bound to all synapses such that *w*_*i*_ is set to zero if *w*_*i*_ crosses zero from above.

Parameters for the plasticity model were manually selected so as to replicate classic spike-timing dependent plasticity experiments (Supplemental Figure S1) and to qualitatively reproduce the effects of DCS on LTP. We are mainly concerned with relative changes in LTP due to DCS (or spike timing/frequency in the case of replicating STDP experiments) and so do not adjust parameters to quantitatively reproduce the amount of LTP in each experiment. Under these constraints we were able to use the same set of parameters for each simulation (Supplemental Table 1). Numerical integration using the forward euler method (0.025 ms time step) was used to solve for *w*_*i*_*(t)*

### Additional simulation details for Figure 6

To emulate the two-pathway experiments of Figure 3, a population of cells was generated as described above, but now two groups of synapses were selected to be activated on each cell, a strong and a weak group. Note that because synapses were selected randomly with replacement, a given synapse was allowed to be part of both groups, although this was rare. As in our experiments, three sets of simulations were run: activation of only the weak pathway at 5 Hz, activation of only the strong pathway with TBS, or activation of both pathways simultaneously.

We hypothesized that pairing the two pathways boosted LTP by facilitating spikes that backpropagate from the soma to synapses in both pathways. To test this hypothesis in our model we wanted to measure spikes that occurred at each synapse, and importantly whether a given spike was initiated in the soma. Of course, if a spike is initiated in the soma, it should occur before a spike is observed in the dendrite. To evaluate this time difference, we first detected the onset of spikes in each segment of the model neuron by measuring the time at which the voltage crossed a threshold of −30 mV from below.

For each segment, a binary vector of spike times was therefore created, with each entry corresponding to a time step in the simulation (1=spike detected, 0=no spike detected). The cross correlation was computed between this spike vector and the corresponding vector measured at the soma. This yields binary vector for each segment, where each entry corresponds to a possible time delay between that segment and the soma. A value of 1 in this vector indicates that the corresponding delay was observed. By averaging this cross correlation over all activated synaptic locations, we get a probability density over different spike delays between the soma and dendrite. In general, a spike can propagate throughout the entire neuron within ~2 ms. We therefore assume that temporal correlations occurring within this +−2 ms window correspond to delays that are due to the propagation of a single spike, while correlations that are outside of this +−2 ms window correspond to delays between different spikes. We have set up the analysis so negative time delays correspond to spikes that appeared in the soma first. Spikes that initiate in the soma and propagate to the dendrite should therefore add density between −2 and 0 ms (Figure 6B).

The metric based on spike cross-correlation only captures spike events that occur in both the soma and dendrite. However, spikes can also initiate in the dendrite, but may not propagate completely to the soma. These local spikes would also make a large contribution to synaptic plasticity at a subset of local synapses, but do not contribute to the cross correlation metric. We therefore also considered the overall number of dendritic spikes (global and local) as a function of time during each theta burst at which they occurred. We divided the simulation into individual theta bursts, and within each burst, the simulation was divided into 1 ms time bins. Spikes were counted in these time bins across all synapses. By summing across all synapses, we get the total number of dendritic spikes that occur as a function of the time since burst onset (Figure 6C).

### Simulation details for Figure 8

Membrane polarization (Figure 8A,B) was calculated by simulating a single cell without synaptic input for 100 ms with varying applied electric field. Membrane polarization due to DCS was calculated in each compartment as the voltage at the end of the simulation minus the corresponding voltage in the control simulation without DCS.

For each simulation and each activated synapse *k*, we calculate the effect of DCS on plasticity

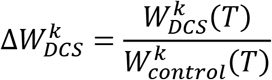

where *T* is the duration of the simulation, 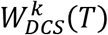 is the final weight of the *k*^*th*^ synapse at the end of the simulation with DCS, 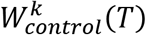 is the weight at the end of the corresponding control simulation where no DCS was applied. Note that all DCS simulations have a control simulation in which all other details are identical. Therefore, any deviation of 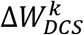 from 1 represents the effect of DCS on the *k*^*th*^ synaptic weight.

For a given DCS waveform (polarity and magnitude), we are interested in the distribution of 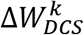 over all *k* synapses in the population. Figure 8E displays the mean of this distribution as a function of DCS intensity.

## Abbreviations

tDCS: transcranial direct current stimulation
DCS: direct current stimulation
LTP: long term potentiation
TBS: theta burst stimulation
STDP: spike-timing dependent plasticity

## Author contributions

G.K. and A.R. designed and executed experiments, M.S. executed experiments, G.K. designed and ran simulations, G.K. analyzed data, G.K., A.R., M.B., L.P. interpreted results and wrote the manuscript.

## Funding sources

This work was supported by the National Institutes of Health (grant number 5R01NS095123)

## Competing interests

LCP and MB have shares in Soterix Medical Inc and are listed as inventors in patents of the City University of New York related to high-definition tDCS.

## Supplemental Material

### Simulation of STDP experiments

To help constrain our computational model, we simulated canonical STDP results in the literature (26,43). First, we simulated STDP by pairing spiking generated at the soma with synaptic inputs on the proximal apical dendrites (5 synapses, randomly distributed). Somatic spikes were evoked by a 5 ms, 1 nA current injection in the soma at varying temporal offsets from synaptic input (Δt), with positive Δt corresponding to pre before post pairing (pre-post) and negative Δt corresponding to post before pre pairing (post-pre). Synaptic weights at the end of the simulation were normalized to the initial baseline value and plotted as a function of Δt (Figure S1A). The detailed neuron model with the specified plasticity parameters (Table 1) qualitatively reproduces the canonical STDP window (Figure S1A), where pre-post pairing leads to potentiation and post-pre pairing leads to depression. We next simulated the experimentally observed frequency-dependence of STDP (26,43). Here we performed similar simulations but with Δt fixed at either −10 or +10 ms and varied the frequency of pre and postsynaptic pairings (1, 5, 10, 20, 30, 40, 50, 75, 100 Hz, Figure S1B).

**Figure S1.**
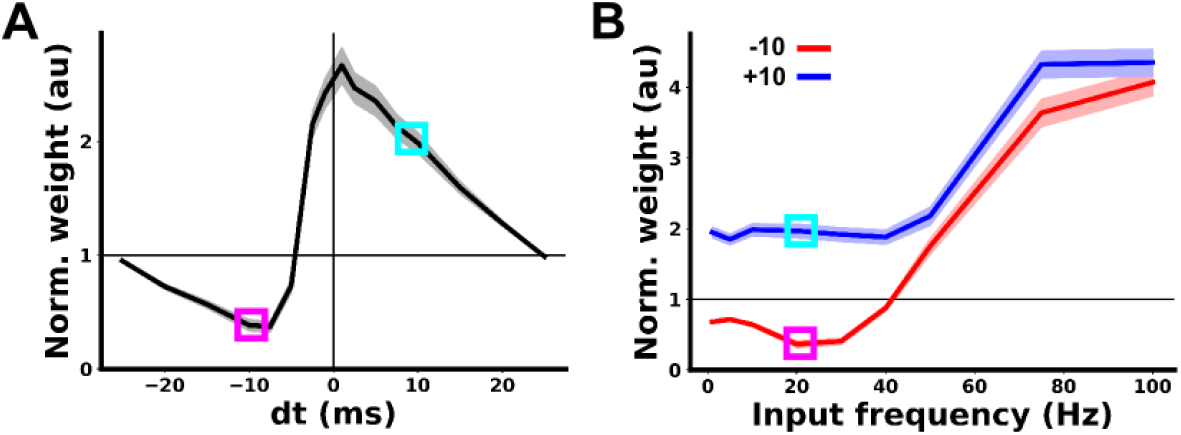
Model reproduces classic stdp with frequency dependence. **A)** Final synaptic weight (average across the entire population of synapses) as a function of pre-post timing for 6 pairings at 20 Hz. Positive dt corresponds to pre-post pairings, while negative dt corresponds to post-pre pairings. **B)** Final synaptic weight (average across the entire population of synapses) as a function of pairing frequency in STDP simulations. The red curve corresponds to 6 post-pre pairings (Δt =−10 ms). The blue curve corresponds to 6 pre-post pairings (Δt =+10 ms). The cyan and magenta boxes mark data points that are from identical simulation in A and B

**Figure S2.**
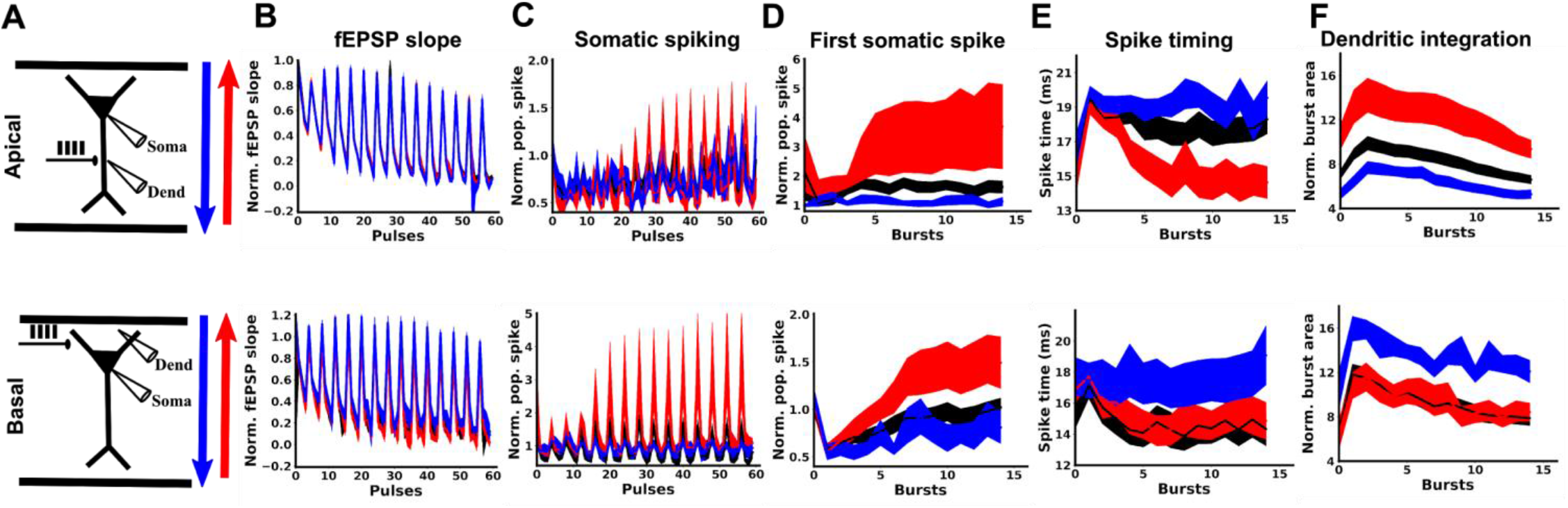
Extracellular voltage dynamics during induction. **A)** Schematic of experimental design for TBS experiments in apical (top row) and basal (bottom row) dendrites, depicting the orientation of anodal (red) and cathodal (blue) electric fields with respect to CA1 pyramidal cells. Black traces indicate control experiments, where no electric field was applied. **B)** DCS has no significant effect on fEPSP slopes recorded during induction. **C)** Anodal DCS enhances population spikes recorded at the soma in response to both apical and basal synaptic activity. **D)** Same data as in C, but showing on the first pulse during each burst of the TBS protocol. The effect of DCS is most pronounced on the first pulse. **E)** DCS shifts average spike timing for each burst during induction (see methods “quantifying somatic activity” for details) **F)** DCS has opposite effects on dendritic integration in response to apical or basal synaptic input. The horizontal axes represent either the number of individual bipolar stimulus pulses (60 in total) or bursts (15 in total) applied to activate synapses during induction. All data normalized to the mean of the 20 baseline responses before induction and are represented as mean±s.e.m.

**Figure S3.**
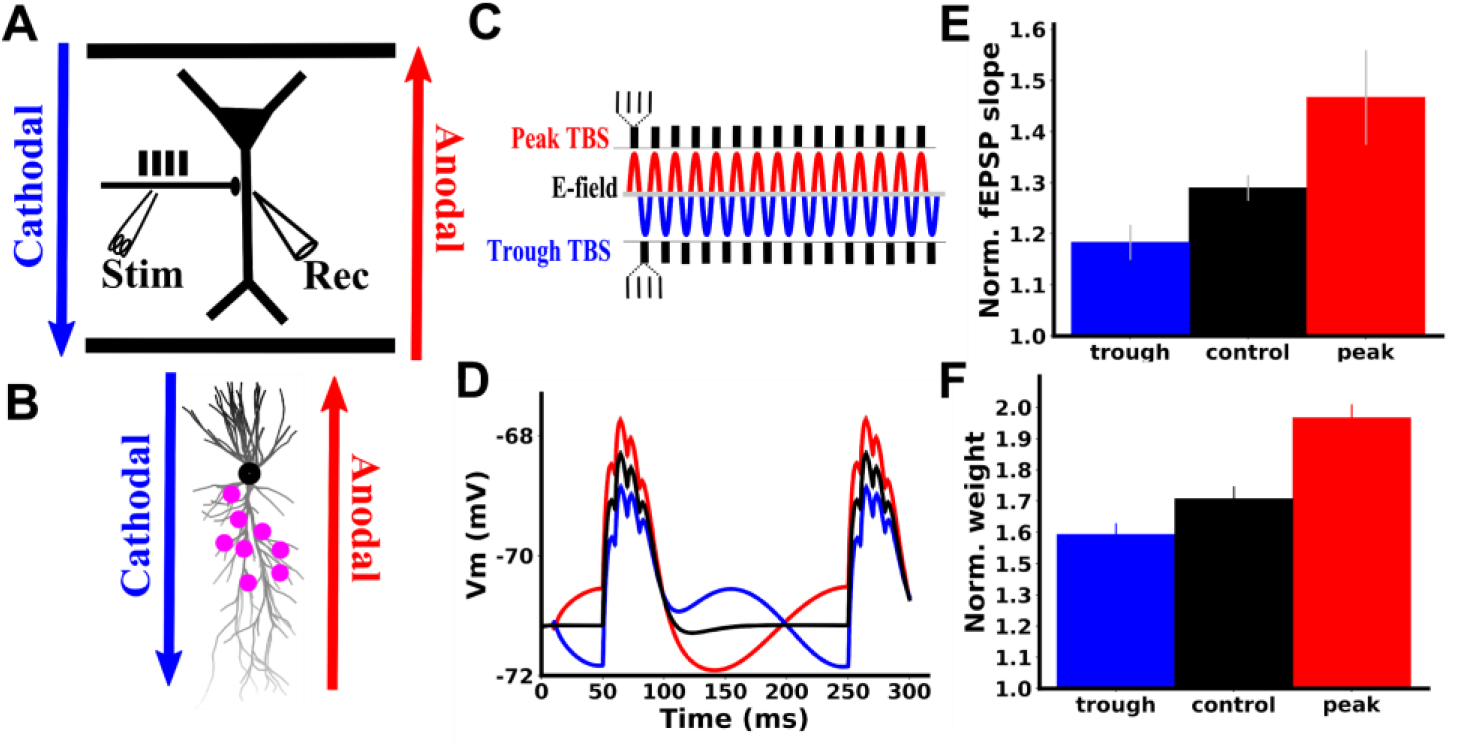
Model reproduces effects of AC stimulation. **A)** Schematic of experimental design (top) and model neuron morphology and synapse distribution (bottom). **B)** Timing of synaptic inputs and applied electric field for both experiment and model. **C)** For peak TBS (red), each burst during the TBS protocol is timed to the peak of the extracellular field, such that pyramidal cell somas are depolarized when the synaptic inputs arrive. For trough TBS (blue), each burst during the TBS protocol is timed to the trough of the extracellular field, such that pyramidal cell somas are hyperpolarized when the synaptic inputs arrive. **D)** Example voltage traces from somatic compartment of model neuron during first two bursts of simulation. **E)** Resulting experimental LTP in each condition. As in Figure 1C, fEPSP slopes are averaged over the last 10 minutes of recording in each condition. **F)** Model LTP predictions qualitatively match (same direction of DCS effect) experimental LTP results (D). The vertical axis (Norm. weight) is the average weight of all weak pathway synapses at the end of simulation, calculated offline using the learning rule (41).

**Figure S4.**
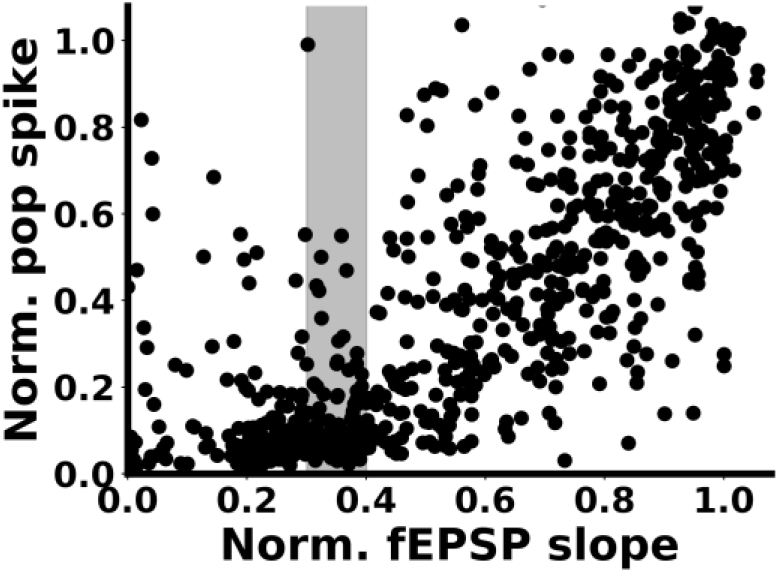
Input-output curve reveals that the baseline of our experiments is set near a nonlinearity. Baseline population spike amplitude as a function of baseline fEPSP slope for all slices. fEPSP slopes are normalized to the maximum value detected in the process of setting baseline bipolar stimulus intensity (see methods “fEPSP recordings”). The horizontal axis can therefore be thought of as the fraction of activated synapses in the population. Population spikes are normalized to the population spike magnitude recorded when the maximum fEPSP is established. The gray box highlights approximately where baseline fEPSP’s were set before running LTP experiments (30-40% of maximum). Note that experiments are run near a nonlinearity in the input-output curve, such that system is more responsive to increases in input rather than decreases in input.

**Figure S5.**
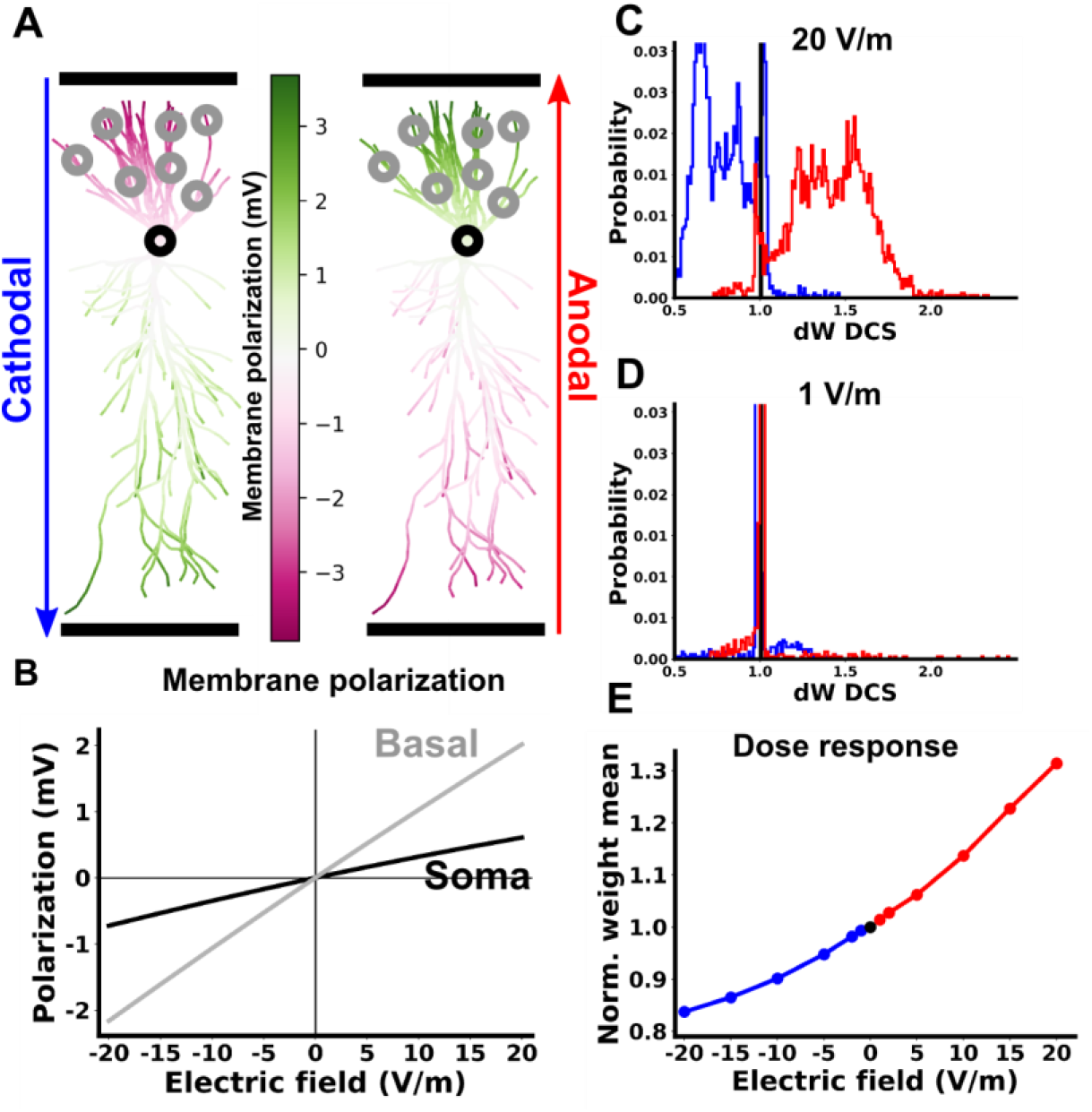
Dose response in computational model for TBS in basal dendrites. Same as Figure 8 but for basal dendrites.

### Acute effects on synaptic input

To rule out potential effects of DCS directly on the recruitment of presynaptic axons, we analyzed acute effects of DCS on fiber volleys, acute fEPSP slope, and paired-pulse ratio (PPR). To capture potential acute effects of DCS, all data are taken from the first pulse during induction (first two pulses for PPR), and normalized to the mean of baseline responses. Fiber volleys were calculated by measuring the dendritic extracellular voltage at 1 ms and 2 ms after bipolar pulse onset. A line was fit between these two points, and the unnormalized fiber volleys were taken as the average voltage below this line. Paired pulse ratio is taken as the ratio of the second to the first normalized fEPSP slope during TBS induction (10 ms inter-pulse interval).

We found no significant effects of DCS on fEPSP slope (Figure S6A; control 1.006±0.007, N=31; anodal 0.98±0.016, N=21, p=0.14 vs. control, cathodal 1.017±0.014, N=11, p=0.505 vs. control), fiber volleys (Figure S6B; control 1.012±0.038, N=28; anodal 1.039±0.03, N=13, p=0.65 vs. control; cathodal 1.007±0.084, N=3, p=0.966 vs. control), or paired pulse ratio (Figure S6C; control 0.766±0.022, N=31; anodal 0.758±0.022, N=21, p=0.816 vs. control; cathodal 0.703±0.033, N=11, p=0.166 vs. control). We note that PPR is typically measured with longer inter-pulse interval (e.g. 50 ms). However, if presynaptic release is altered by DCS, then this should be reflected in PPR measured with the interval used here. Moreover, our previous study with an identical setup (14) found no effects on PPR with a 50 ms interval.

**Figure S6.**
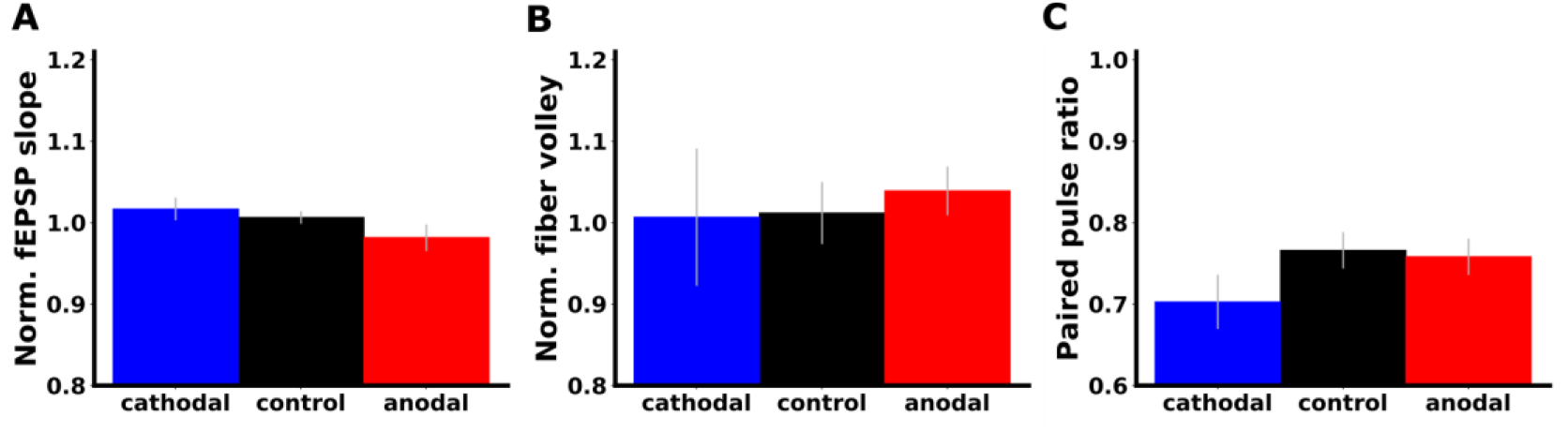
Acute effects of DCS on measures of presynaptic release. DCS did not have significant effects on fEPSP slope (**A**), fiber volleys (**B**), or paired pulse ratio (**C**).

**Supplemental Table 1.** Parameters for voltage-based plasticity rule.

| Parameter | Value | Explanation |
| --- | --- | --- |
| $A_{LTD}$ | $0.1 \text{ mV}^{-1}$ | LTD rate |
| $A_{LTP}$ | $0.04 \text{ mV}^{-2}$ | LTP rate |
| $\theta_-$ | -70 mV | LTD threshold |
| $\theta_+$ | -67 mV | LTP threshold |
| $\tau_x$ | 8 ms | presynaptic input trace lowpass time constant |
| $\tau_-$ | 20 ms | LTD voltage trace lowpass time constant |
| $\tau_+$ | 3 ms | LTP voltage trace lowpass time constant |

**Supplemental Table 2.** Neuron model synaptic parameters.

| Parameter | Value | Explanation |
| --- | --- | --- |
| $\tau_{ampa}^{open}$ | 0.2 ms | AMPA receptor conductance rise time constant |
| $\tau_{ampa}^{close}$ | 2 ms | AMPA receptor conductance decay time constant |
| $\tau_{nmda}^{open}$ | 1 ms | NMDA receptor conductance rise time constant |
| $\tau_{nmda}^{close}$ | 50 ms | NMDA receptor conductance decay time constant |
| $g_{ampa}$ | 1 nS | peak AMPA receptor conductance |
| $g_{nmda}$ | 1 nS | peak NMDA receptor conductance |
| $\tau_F$ | 94 ms | Facilitation time constant |
| $\tau_{D_1}$ | 540 ms | 1st depression time constant |
| $\tau_{D_2}$ | 45 ms | 2nd depression time constant |
| $\tau_{D_3}$ | 120 s | 3rd depression time constant |
| $f$ | 5 | Additive facilitation factor |
| $d_1$ | 0.45 | 1st Multiplicative depression factor |
| $d_2$ | 0.12 | 2nd Multiplicative depression factor |
| $d_3$ | 0.98 | 3rd Multiplicative depression factor |

**Supplemental Table 3.** Neuron model membrane conductance parameters.

| Parameter | Value | Explanation |
| --- | --- | --- |
| $g_{Na^+}^{dend}$ | 25 mS*cm<sup>-2</sup> | Voltage gated sodium conductance in dendrites |
| $g_{Na^+}^{soma}$ | 37.5 mS*cm<sup>-2</sup> | Voltage gated sodium conductance in soma |
| $g_{Na^+}^{axon}$ | 2500 mS*cm<sup>-2</sup> | Voltage gated sodium conductance in axon |
| $E_{Na^+}$ | 55 mV | Sodium reversal potential |
| $E_{K^+}$ | -90 mV | Potassium reversal potential |
| $E_h$ | -30 mV | H-current reversal potential |
| $E_{Ca^{2+}}$ | 140 mV | Calcium reversal potential |
| $g_{K_{DR}^+}$ | 10 mS*cm<sup>-2</sup> | Delayed rectifier potassium peak conductance |
| $g_{K_A^+}$ | 30 mS*cm<sup>-2</sup> | A-type potassium peak conductance |
| $g_h$ | .05*(1+3d/100)<br>mS*cm<sup>-2</sup> | H-channel conductance. Linearly increasing with distance d (in μm) from the soma |
| $g_{Ca_{Lv}^{2+}}$ | 1.25 mS*cm<sup>-2</sup> | L-type calcium channel peak conductance |
| $V_{1/2}^{hprox}$ | -82 mV | Activation threshold for proximal (<100 μm from soma) h channel conductance |
| $V_{1/2}^{hdist}$ | -90 mV | Activation threshold for distal (>100 μm from soma) h channel conductance |
| $\tau_{ampa}^{close}$ | 2 ms | AMPA receptor conductance decay time constant |
| $\tau_{nmda}^{open}$ | 1 ms | NMDA receptor conductance rise time constant |
| $\tau_{nmda}^{close}$ | 50 ms | NMDA receptor conductance decay time constant |
| $g_{ampa}$ | 1 nS | peak AMPA receptor conductance |
| $g_{nmda}$ | 1 nS | peak NMDA receptor conductance |

## References

1. Mancuso LE, Ilieva IP, Hamilton RH, Farah MJ. Does Transcranial Direct Current Stimulation Improve Healthy Working Memory?: A Meta-analytic Review. J Cogn Neurosci. 2016 Apr 7;28(8):1063–89.

2. Kekic M, Boysen E, Campbell IC, Schmidt U. A systematic review of the clinical efficacy of transcranial direct current stimulation (tDCS) in psychiatric disorders. J Psychiatr Res. 2016 Mar 1;74:70–86.

3. Dedoncker J, Brunoni AR, Baeken C, Vanderhasselt M-A. A Systematic Review and Meta-Analysis of the Effects of Transcranial Direct Current Stimulation (tDCS) Over the Dorsolateral Prefrontal Cortex in Healthy and Neuropsychiatric Samples: Influence of Stimulation Parameters. Brain Stimulat. 2016 Jul 1;9(4):501–17.

4. Tedesco Triccas L, Burridge JH, Hughes AM, Pickering RM, Desikan M, Rothwell JC, et al. Multiple sessions of transcranial direct current stimulation and upper extremity rehabilitation in stroke: A review and meta-analysis. Clin Neurophysiol. 2016 Jan 1;127(1):946–55.

5. Santarnecchi E, Brem A-K, Levenbaum E, Thompson T, Kadosh RC, Pascual-Leone A. Enhancing cognition using transcranial electrical stimulation. Curr Opin Behav Sci. 2015 Aug 1;4:171–8.

6. Stagg CJ, Nitsche MA. Physiological Basis of Transcranial Direct Current Stimulation. The Neuroscientist. 2011 Feb 1;17(1):37–53.

7. Gellner A-K, Reis J, Fritsch B. Glia: A Neglected Player in Non-invasive Direct Current Brain Stimulation. Front Cell Neurosci [Internet]. 2016 [cited 2019 Feb 26];10. Available from: https://www.frontiersin.org/articles/10.3389/fncel.2016.00188/full

8. Pelletier SJ, Cicchetti F. Cellular and Molecular Mechanisms of Action of Transcranial Direct Current Stimulation: Evidence from In Vitro and In Vivo Models. Int J Neuropsychopharmacol. 2015 Jan 1;18(2):pyu047–pyu047.

9. Monai H, Ohkura M, Tanaka M, Oe Y, Konno A, Hirai H, et al. Calcium imaging reveals glial involvement in transcranial direct current stimulation-induced plasticity in mouse brain. Nat Commun. 2016 Mar 22;7:11100.

10. Nitsche MA, Fricke K, Henschke U, Schlitterlau A, Liebetanz D, Lang N, et al. Pharmacological Modulation of Cortical Excitability Shifts Induced by Transcranial Direct Current Stimulation in Humans. J Physiol. 2003 Nov;553(1):293–301.

11. Marquez-Ruiz J, Leal-Campanario R, Sanchez-Campusano R, Molaee-Ardekani B, Wendling F, Miranda PC, et al. Transcranial direct-current stimulation modulates synaptic mechanisms involved in associative learning in behaving rabbits. Proc Natl Acad Sci. 2012 Apr 24;109(17):6710–5.

12. Podda MV, Cocco S, Mastrodonato A, Fusco S, Leone L, Barbati SA, et al. Anodal transcranial direct current stimulation boosts synaptic plasticity and memory in mice via epigenetic regulation of Bdnf expression. Sci Rep. 2016 Feb 24;6:22180.

13. Fritsch B, Reis J, Martinowich K, Schambra HM, Ji Y, Cohen LG, et al. Direct Current Stimulation Promotes BDNF-Dependent Synaptic Plasticity: Potential Implications for Motor Learning. Neuron. 2010 Apr;66(2):198–204.

14. Kronberg G, Bridi M, Abel T, Bikson M, Parra LC. Direct Current Stimulation Modulates LTP and LTD: Activity Dependence and Dendritic Effects. Brain Stimulat. 2017 Jan;10(1):51–8.

15. Ranieri F, Podda MV, Riccardi E, Frisullo G, Dileone M, Profice P, et al. Modulation of LTP at rat hippocampal CA3-CA1 synapses by direct current stimulation. J Neurophysiol. 2012 Apr;107(7):1868–80.

16. Sun Y, Lipton JO, Boyle LM, Madsen JR, Goldenberg MC, Pascual-Leone A, et al. Direct current stimulation induces mGluR5-dependent neocortical plasticity: Cathodal DCS Induces LTD. Ann Neurol. 2016 Aug;80(2):233–46.

17. Akiyama H, Shimizu Y, Miyakawa H, Inoue M. Extracellular DC electric fields induce nonuniform membrane polarization in rat hippocampal CA1 pyramidal neurons. Brain Res. 2011 Apr 6;1383:22–35.

18. Bikson M, Inoue M, Akiyama H, Deans JK, Fox JE, Miyakawa H, et al. Effects of uniform extracellular DC electric fields on excitability in rat hippocampal slices *in vitro*: Modulation of neuronal function by electric fields. J Physiol. 2004 May;557(1):175–90.

19. Radman T, Ramos RL, Brumberg JC, Bikson M. Role of cortical cell type and morphology in subthreshold and suprathreshold uniform electric field stimulation in vitro. Brain Stimulat. 2009 Oct;2(4):215–28, 228.e1-3.

20. Deans JK, Powell AD, Jefferys JGR. Sensitivity of coherent oscillations in rat hippocampus to AC electric fields. J Physiol. 2007 Sep 1;583(Pt 2):555–65.

21. Chakraborty D, Truong DQ, Bikson M, Kaphzan H. Neuromodulation of Axon Terminals. Cereb Cortex. 2018 Aug 1;28(8):2786–94.

22. Rahman A, Reato D, Arlotti M, Gasca F, Datta A, Parra LC, et al. Cellular effects of acute direct current stimulation: somatic and synaptic terminal effects. J Physiol. 2013 May 15;591(Pt 10):2563–78.

23. Aspart F, Remme MWH, Obermayer K. Differential polarization of cortical pyramidal neuron dendrites through weak extracellular fields. Graham LJ, editor. PLOS Comput Biol. 2018 May 4;14(5):e1006124.

24. Artola A, Bröcher S, Singer W. Different voltage-dependent thresholds for inducing long-term depression and long-term potentiation in slices of rat visual cortex. Nature. 1990 Sep 6;347(6288):69–72.

25. Ngezahayo A, Schachner M, Artola A. Synaptic activity modulates the induction of bidirectional synaptic changes in adult mouse hippocampus. J Neurosci Off J Soc Neurosci. 2000 Apr 1;20(7):2451–8.

26. Sjöström PJ, Turrigiano GG, Nelson SB. Rate, Timing, and Cooperativity Jointly Determine Cortical Synaptic Plasticity. Neuron. 2001 Dec;32(6):1149–64.

27. Sjöström PJ, Häusser M. A Cooperative Switch Determines the Sign of Synaptic Plasticity in Distal Dendrites of Neocortical Pyramidal Neurons. Neuron. 2006 Jul;51(2):227–38.

28. Simonsmeier BA, Grabner RH, Hein J, Krenz U, Schneider M. Electrical brain stimulation (tES) improves learning more than performance: A meta-analysis. Neurosci Biobehav Rev. 2018 Jan;84:171–81.

29. Horvath JC, Forte JD, Carter O. Quantitative Review Finds No Evidence of Cognitive Effects in Healthy Populations From Single-session Transcranial Direct Current Stimulation (tDCS). Brain Stimulat. 2015 May;8(3):535–50.

30. Horvath JC, Forte JD, Carter O. Evidence that transcranial direct current stimulation (tDCS) generates little-to-no reliable neurophysiologic effect beyond MEP amplitude modulation in healthy human subjects: A systematic review. Neuropsychologia. 2015 Jan;66:213–36.

31. Hashemirad F, Zoghi M, Fitzgerald PB, Jaberzadeh S. The effect of anodal transcranial direct current stimulation on motor sequence learning in healthy individuals: A systematic review and meta-analysis. Brain Cogn. 2016 Feb;102:1–12.

32. Cabral ME, Baltar A, Borba R, Galvão S, Santos L, Fregni F, et al. Transcranial direct current stimulation: before, during, or after motor training? NeuroReport. 2015 Aug;26(11):618–22.

33. Besson P, Muthalib M, Dray G, Rothwell J, Perrey S. Concurrent anodal transcranial direct-current stimulation and motor task to influence sensorimotor cortex activation. Brain Res. 2019 May;1710:181–7.

34. Stagg CJ, Jayaram G, Pastor D, Kincses ZT, Matthews PM, Johansen-Berg H. Polarity and timing-dependent effects of transcranial direct current stimulation in explicit motor learning. Neuropsychologia. 2011 Apr;49(5):800–4.

35. Bliss TVP, Collingridge GL. A synaptic model of memory: long-term potentiation in the hippocampus. Nature. 1993 Jan;361(6407):31–9.

36. Litwin-Kumar A, Doiron B. Formation and maintenance of neuronal assemblies through synaptic plasticity. Nat Commun [Internet]. 2014 Dec [cited 2019 Feb 27];5(1). Available from: http://www.nature.com/articles/ncomms6319

37. Holtmaat A, Caroni P. Functional and structural underpinnings of neuronal assembly formation in learning. Nat Neurosci. 2016 Dec;19(12):1553–62.

38. Migliore M, Ferrante M, Ascoli GA. Signal Propagation in Oblique Dendrites of CA1 Pyramidal Cells. J Neurophysiol. 2005 Dec;94(6):4145–55.

39. Migliore M, Hoffman DA, Magee JC, Johnston D. Role of an A-type K+ conductance in the back-propagation of action potentials in the dendrites of hippocampal pyramidal neurons. J Comput Neurosci. 1999 Aug;7(1):5–15.

40. Hines ML, Carnevale NT. The NEURON Simulation Environment. Neural Comput. 1997 Aug;9(6):1179–209.

41. Clopath C, Büsing L, Vasilaki E, Gerstner W. Connectivity reflects coding: a model of voltage-based STDP with homeostasis. Nat Neurosci. 2010 Mar;13(3):344–52.

42. Ko H, Cossell L, Baragli C, Antolik J, Clopath C, Hofer SB, et al. The emergence of functional microcircuits in visual cortex. Nature. 2013 Apr;496(7443):96–100.

43. Bono J, Clopath C. Modeling somatic and dendritic spike mediated plasticity at the single neuron and network level. Nat Commun [Internet]. 2017 Dec [cited 2019 Jun 3];8(1). Available from: http://www.nature.com/articles/s41467-017-00740-z

44. Golding NL, Staff NP, Spruston N. Dendritic spikes as a mechanism for cooperative long-term potentiation. Nature. 2002 Jul 18;418(6895):326–31.

45. Huang Y, Liu AA, Lafon B, Friedman D, Dayan M, Wang X, et al. Measurements and models of electric fields in the in vivo human brain during transcranial electric stimulation. eLife [Internet]. 2017 Feb 7 [cited 2019 Jun 12];6. Available from: https://elifesciences.org/articles/18834

46. Opitz A, Falchier A, Yan C-G, Yeagle EM, Linn GS, Megevand P, et al. Spatiotemporal structure of intracranial electric fields induced by transcranial electric stimulation in humans and nonhuman primates. Sci Rep [Internet]. 2016 Aug [cited 2019 Jun 12];6(1). Available from: http://www.nature.com/articles/srep31236

47. Cao G, Harris KM. Augmenting saturated LTP by broadly spaced episodes of theta-burst stimulation in hippocampal area CA1 of adult rats and mice. J Neurophysiol. 2014 Oct 15;112(8):1916–24.

48. Nitsche MA, Fricke K, Henschke U, Schlitterlau A, Liebetanz D, Lang N, et al. Pharmacological modulation of cortical excitability shifts induced by transcranial direct current stimulation in humans. J Physiol. 2003 Nov 15;553(Pt 1):293–301.

49. Hebb DO. The Organization of Behavior: A Neuropsychological Theory. Psychology Press; 2005. 379 p.

50. Abbott LF, Nelson SB. Synaptic plasticity: taming the beast. Nat Neurosci. 2000 Nov 1;3(11s):1178–83.

51. Larson J, Munkácsy E. Theta-burst LTP. Brain Res. 2015 Sep;1621:38–50.

52. Palm G. Neural associative memories and sparse coding. Neural Netw. 2013 Jan;37:165–71.

53. Malenka RC, Bear MF. LTP and LTD. Neuron. 2004 Sep;44(1):5–21.

54. Hamoudi M, Schambra HM, Fritsch B, Schoechlin-Marx A, Weiller C, Cohen LG, et al. Transcranial Direct Current Stimulation Enhances Motor Skill Learning but Not Generalization in Chronic Stroke. Neurorehabil Neural Repair. 2018 Apr;32(4-5):295–308.

55. Pisoni A, Mattavelli G, Papagno C, Rosanova M, Casali AG, Romero Lauro LJ. Cognitive Enhancement Induced by Anodal tDCS Drives Circuit-Specific Cortical Plasticity. Cereb Cortex. 2018 Apr 1;28(4):1132–40.

56. Martin DM, Liu R, Alonzo A, Green M, Loo CK. Use of transcranial direct current stimulation (tDCS) to enhance cognitive training: effect of timing of stimulation. Exp Brain Res. 2014 Oct;232(10):3345–51.

57. Rioult-Pedotti M-S. Learning-Induced LTP in Neocortex. Science. 2000 Oct 20;290(5491):533–6.

58. Cichon J, Gan W-B. Branch-specific dendritic Ca2+ spikes cause persistent synaptic plasticity. Nature. 2015 Apr;520(7546):180–5.

59. Ziemann U. Learning Modifies Subsequent Induction of Long-Term Potentiation-Like and Long-Term Depression-Like Plasticity in Human Motor Cortex. J Neurosci. 2004 Feb 18;24(7):1666–72.

60. Citri A, Malenka RC. Synaptic Plasticity: Multiple Forms, Functions and Mechanisms. Neuropsychopharmacology. 2008 Jan;33(1):18–41.

61. Lafon B, Rahman A, Bikson M, Parra LC. Direct Current Stimulation Alters Neuronal Input/Output Function. Brain Stimulat. 2017 Feb;10(1):36–45.

62. Nitsche MA, Schauenburg A, Lang N, Liebetanz D, Exner C, Paulus W, et al. Facilitation of Implicit Motor Learning by Weak Transcranial Direct Current Stimulation of the Primary Motor Cortex in the Human. J Cogn Neurosci. 2003 May;15(4):619–26.

63. Kincses TZ, Antal A, Nitsche MA, Bártfai O, Paulus W. Facilitation of probabilistic classification learning by transcranial direct current stimulation of the prefrontal cortex in the human. Neuropsychologia. 2004;42(1):113–7.

64. Migliore R, De Simone G, Migliore M. Induction of long-term potentiation and depression in individual synapses of CA1 pyramidal neurons. BMC Neurosci [Internet]. 2015 Dec [cited 2019 Jun 14];16(S1). Available from: https://bmcneurosci.biomedcentral.com/articles/10.1186/1471-2202-16-S1-P164

65. Reato D, Rahman A, Bikson M, Parra LC. Low-intensity electrical stimulation affects network dynamics by modulating population rate and spike timing. J Neurosci Off J Soc Neurosci. 2010 Nov 10;30(45):15067–79.

66. Terzuolo CA, Bullock TH. MEASUREMENT OF IMPOSED VOLTAGE GRADIENT ADEQUATE TO MODULATE NEURONAL FIRING. Proc Natl Acad Sci U S A. 1956 Sep;42(9):687–94.

67. Bindman LJ, Lippold OC, Redfearn JW. the action of brief polarizing currents on the cerebral cortex of the rat (1) during current flow and (2) in the production of long-lasting after-effects. J Physiol. 1964 Aug;172:369–82.

68. Chan CY, Hounsgaard J, Nicholson C. Effects of electric fields on transmembrane potential and excitability of turtle cerebellar Purkinje cells in vitro. J Physiol. 1988 Aug;402:751–71.

69. Radman T, Su Y, An JH, Parra LC, Bikson M. Spike timing amplifies the effect of electric fields on neurons: implications for endogenous field effects. J Neurosci Off J Soc Neurosci. 2007 Mar 14;27(11):3030–6.

70. Liu A, Vöröslakos M, Kronberg G, Henin S, Krause MR, Huang Y, et al. Immediate neurophysiological effects of transcranial electrical stimulation. Nat Commun [Internet]. 2018 Dec [cited 2019 Feb 27];9(1). Available from: http://www.nature.com/articles/s41467-018-07233-7

71. Nitsche MA, Paulus W. Excitability changes induced in the human motor cortex by weak transcranial direct current stimulation. J Physiol. 2000 Sep 15;527 Pt 3:633–9.

72. Nitsche MA, Paulus W. Sustained excitability elevations induced by transcranial DC motor cortex stimulation in humans. Neurology. 2001 Nov 27;57(10):1899–901.

73. Monai H, Ohkura M, Tanaka M, Oe Y, Konno A, Hirai H, et al. Calcium imaging reveals glial involvement in transcranial direct current stimulation-induced plasticity in mouse brain. Nat Commun. 2016 Mar 22;7:11100.

74. Bachtiar V, Near J, Johansen-Berg H, Stagg CJ. Modulation of GABA and resting state functional connectivity by transcranial direct current stimulation. eLife [Internet]. 2015 Sep 18 [cited 2019 Jun 17];4. Available from: https://elifesciences.org/articles/08789

75. Paciello F, Podda MV, Rolesi R, Cocco S, Petrosini L, Troiani D, et al. Anodal transcranial direct current stimulation affects auditory cortex plasticity in normal-hearing and noise-exposed rats. Brain Stimulat. 2018 Sep;11(5):1008–23.

76. Edelmann E, Lessmann V, Brigadski T. Pre- and postsynaptic twists in BDNF secretion and action in synaptic plasticity. Neuropharmacology. 2014 Jan;76 Pt C:610–27.

77. Edelmann E, Cepeda-Prado E, Franck M, Lichtenecker P, Brigadski T, Leßmann V. Theta Burst Firing Recruits BDNF Release and Signaling in Postsynaptic CA1 Neurons in Spike-Timing-Dependent LTP. Neuron. 2015 May;86(4):1041–54.

78. Leschik J, Eckenstaler R, Endres T, Munsch T, Edelmann E, Richter K, et al. Prominent Postsynaptic and Dendritic Exocytosis of Endogenous BDNF Vesicles in BDNF-GFP Knock-in Mice. Mol Neurobiol [Internet]. 2019 Mar 30 [cited 2019 Jun 14]; Available from: http://link.springer.com/10.1007/s12035-019-1551-0

79. Hoare JI, Rajnicek AM, McCaig CD, Barker RN, Wilson HM. Electric fields are novel determinants of human macrophage functions. J Leukoc Biol. 2016 Jun;99(6):1141–51.

80. Rajnicek A, Gow N, McCaig C. Electric field-induced orientation of rat hippocampal neurones in vitro. Exp Physiol. 1992 Jan 1;77(1):229–32.

81. Mishima T, Nagai T, Yahagi K, Akther S, Oe Y, Monai H, et al. Transcranial direct current stimulation (tDCS) induces adrenergic receptor-dependent microglial morphological changes in mice. eNeuro. 2019 Aug 23;ENEURO.0204-19.2019.

82. Huang Y, Parra LC. Can transcranial electric stimulation with multiple electrodes reach deep targets? Brain Stimulat. 2019 Jan;12(1):30–40.

83. Keck T, Toyoizumi T, Chen L, Doiron B, Feldman DE, Fox K, et al. Integrating Hebbian and homeostatic plasticity: the current state of the field and future research directions. Philos Trans R Soc B Biol Sci. 2017 Mar 5;372(1715):20160158.

84. Rohan JG, Carhuatanta KA, McInturf SM, Miklasevich MK, Jankord R. Modulating Hippocampal Plasticity with In Vivo Brain Stimulation. J Neurosci. 2015 Sep 16;35(37):12824–32.

85. Vöröslakos M, Takeuchi Y, Brinyiczki K, Zombori T, Oliva A, Fernández-Ruiz A, et al. Direct effects of transcranial electric stimulation on brain circuits in rats and humans. Nat Commun [Internet]. 2018 Dec [cited 2019 Feb 27];9(1). Available from: http://www.nature.com/articles/s41467-018-02928-3

86. Hayashi-Takagi A, Yagishita S, Nakamura M, Shirai F, Wu YI, Loshbaugh AL, et al. Labelling and optical erasure of synaptic memory traces in the motor cortex. Nature. 2015 Sep;525(7569):333–8.

87. Hines M. NEURON and Python. Front Neuroinformatics [Internet]. 2009 [cited 2019 Jun 3];3. Available from: http://journal.frontiersin.org/article/10.3389/neuro.11.001.2009/abstract

88. Kim Y, Hsu C-L, Cembrowski MS, Mensh BD, Spruston N. Dendritic sodium spikes are required for long-term potentiation at distal synapses on hippocampal pyramidal neurons. eLife [Internet]. 2015 Aug 6 [cited 2019 Jun 3];4. Available from: https://elifesciences.org/articles/06414

89. Kole MHP, Ilschner SU, Kampa BM, Williams SR, Ruben PC, Stuart GJ. Action potential generation requires a high sodium channel density in the axon initial segment. Nat Neurosci. 2008 Feb;11(2):178–86.

90. Maex R, Schutter ED. Synchronization of Golgi and Granule Cell Firing in a Detailed Network Model of the Cerebellar Granule Cell Layer. J Neurophysiol. 1998 Nov;80(5):2521–37.

91. Spruston N, Jonas P, Sakmann B. Dendritic glutamate receptor channels in rat hippocampal CA3 and CA1 pyramidal neurons. J Physiol. 1995 Jan 15;482 (Pt 2):325–52.

92. Varela JA, Sen K, Gibson J, Fost J, Abbott LF, Nelson SB. A Quantitative Description of Short-Term Plasticity at Excitatory Synapses in Layer 2/3 of Rat Primary Visual Cortex. J Neurosci. 1997 Oct 15;17(20):7926–40.

